# Dynamic membrane organization coordinates protein-O-mannosylation and proteostasis in mycobacteria

**DOI:** 10.64898/2026.09.29.755281

**Authors:** Chloé Rivière, Gabriel Publicola, Alexandre Stella, Odile Burlet-Schiltz, Jérôme Nigou, Michel Rivière

**Affiliations:** Université de Toulouse, CNRS, IPBS, Toulouse, France

**Author notes:** To whom correspondence may be addressed: Institut de Pharmacologie et de Biologie Structurale, Universite de Toulouse, CNRS, IPBS, 205 route de Narbonne, BP 64182, F-31077 Toulouse Cedex 4, France.,; ☎.: 33-561-175-558.

## Abstract

Integral membrane enzymes operate within molecular environments, yet how these are organized remains poorly understood. Using M. smegmatis as a mycobacterial model, we define the membrane environment of the protein O-mannosyltransferase MtPMT through stratified interactomics combining affinity purification–mass spectrometry under four stringencies with a persistence score. This approach resolves a persistent membrane-associated core of 108 proteins from more transient layers, revealing a hierarchically organized landscape enriched in Sec protein translocation, protein biogenesis and envelope proteostasis machineries. Reciprocal affinity purification and in vivo FRET identify YajC and FtsH as neighbors of MtPMT, while structural modelling supports integration within a Sec-associated assembly. Genetic disruption of factors reduces O-mannosylation activity, demonstrating dependence on platform integrity. The MtPMT-associated network is remodelled during growth, indicating an adaptive system. Loss of POM compromises envelope permeability and stress resilience, linking O-mannosylation to fitness.

Together, our results establish protein O-mannosylation as an integrated component of a dynamic membrane platform coordinating secretion, membrane protein biogenesis and proteostasis. Our findings support a model in which mycobacterial membranes contain specialized domains dynamically modulated by physiological state.

## Introduction

Cellular functions are orchestrated through dynamic and spatially organized protein interaction networks [1] [2] [3]. Defining the architecture of these networks remains particularly challenging for integral membrane proteins, whose hydrophobic nature complicates their biochemical isolation while often disrupting native molecular associations. This limitation is especially critical for enzymes embedded within the bacterial envelope, where essential processes such as protein secretion, cell envelope biogenesis, and energy metabolism rely on highly coordinated molecular assemblies. Although biochemical studies have provided insights into individual envelope-associated processes, the spatial organization and functional compartmentalization of these machineries within the complex and specific mycobacterial envelope remain poorly understood [4] [5] [6] [7]. Among these processes is the protein O-mannosylation (POM), an evolutionarily conserved membrane associated post-translational glycosylation process catalyzed by protein O-mannosyltransferases (PMTs), which transfer mannose residues from phospho-isoprenoid donors onto serine or threonine residues of acceptor proteins [8] [9]. Although PMT enzymology [10] [11] [12] [13] [14] [15] [16] [17] [18] and substrate repertoires [19] [20] have been increasingly characterized, considerably less is known about the membrane molecular environments within which these enzymes operate.

In eukaryotes, PMTs are essential [21] [22] [23] and functionally linked to protein translocation and quality-control pathways. PMT activity is associated with components of the endoplasmic reticulum translocon [24] [25], and contributes to unfolded protein O-mannosylation (UPOM), a process coupling glycosylation to protein quality control [26] [27] [28] [29]. These observations suggest functional coordination between PMT activity, protein translocation and proteostasis, although whether such relationships rely on stable or spatially organized molecular assemblies remains unknow.

In bacteria, and particularly in the human pathogen *Mycobacterium tuberculosis* (*Mtb*), PMT contributes critically to physiology and virulence [30] [31] [32] [33]. Multiple *Mtb* mannoproteins participate in host–pathogen interactions, immune modulation and maintenance of cell envelope integrity [20] [34] [35] [36] [37] [38] [39]. Consistent with these functions, the vast majority of characterized mannoproteins are either associated with the mycobacterial cell envelope or secreted into the extracellular environment [20], placing them at the host–pathogen interface. As a result, many of these proteins are exported through the Sec secretion pathway before reaching their final destination [17] [40] [41]. Furthermore, several Sec-exported mannoproteins undergo additional lipidation events, suggesting potential coordination between glycosylation, secretion and lipoprotein maturation [17] [40] [41]. Yet despite these functional connections, the molecular organization surrounding bacterial PMTs and the extent to which envelope-associated pathways may be physically coordinated remain unknown. Comprehensive understanding of the membrane arrangement and mechanistic integration of the proteins involved in this physiological machinery is therefore essential to understand mycobacterial physiology and to identify potential vulnerabilities for therapeutic intervention.

Using functional fusion proteins expressed in the surrogate host *Mycobacterium smegmatis*, we previously characterized the topology and membrane localization [18] of the sole *Mtb* PMT, MtPMT (encoded by *rv1002c,* [17] [42]). MtPMT possesses a compact architecture comprising eleven transmembrane helices and is deeply embedded within the membrane [17] [18], consistent with localization at the interface of envelope-associated processes. However, whether MtPMT functions as an isolated glycosyltransferase or within a broader molecular environment has not been investigated.

Addressing this question remains technically challenging. Integral membrane proteins reside within densely packed and dynamic assemblies, while detergent extraction and biochemical isolation may destabilize weak or transient interactions. Although affinity purification–mass spectrometry (AP–MS) has emerged as a powerful strategy for mapping protein interaction networks [43] [44], resolving the interaction landscape of hydrophobic membrane proteins and distinguishing biologically meaningful partners from membrane-associated background remain major challenges.

To address these limitations, we used an integrated AP–MS framework combining variable reference controls, multiple biochemical stringencies and growth-stage analyses. Beyond individual dataset analysis, we implemented a heuristic “inter-experiment” persistence score to quantify the reproducibility and biochemical stability of MtPMT-associated proteins across distinct purification conditions. By integrating interaction persistence with structural models and functional analyses, we reconstruct the molecular environment of MtPMT and examine whether bacterial protein O-mannosylation is associated with higher-order membrane organization. We further examine the temporal remodeling of this organization by comparing interaction profiles at different stages of bacterial growth

## Results

### Biochemical tuning of AP–MS reveals distinct MtPMT-associated proteomes but a persistent core

To define the molecular environment of the *Mtb* O-mannosyltransferase MtPMT, we employed our previously developed N-terminal mCherry-tagged fusion construct (mC-MtPMT) [18] as bait in affinity purification coupled to mass spectrometry (AP–MS) experiments. This construct, previously validated to retain both proper membrane localization and enzymatic activity [18], was used to capture mC-MtPMT-associated membrane assemblies.

Because the recovery of molecular assemblies is highly sensitive to biochemical conditions— particularly in the case of hydrophobic membrane proteins—we implemented a multi-condition AP–MS strategy designed to explore distinct pre-analytical states of cell lysis, membrane solubilization, and purification stringency. Four independent experimental conditions (C1–C4) were sequentially established by varying specificity controls, detergent solubilization, and ionic strength, with the aim of capturing interacting partners spanning a broad spectrum of biochemical stabilities (**Fig. 1A**).

**Figure 1.**
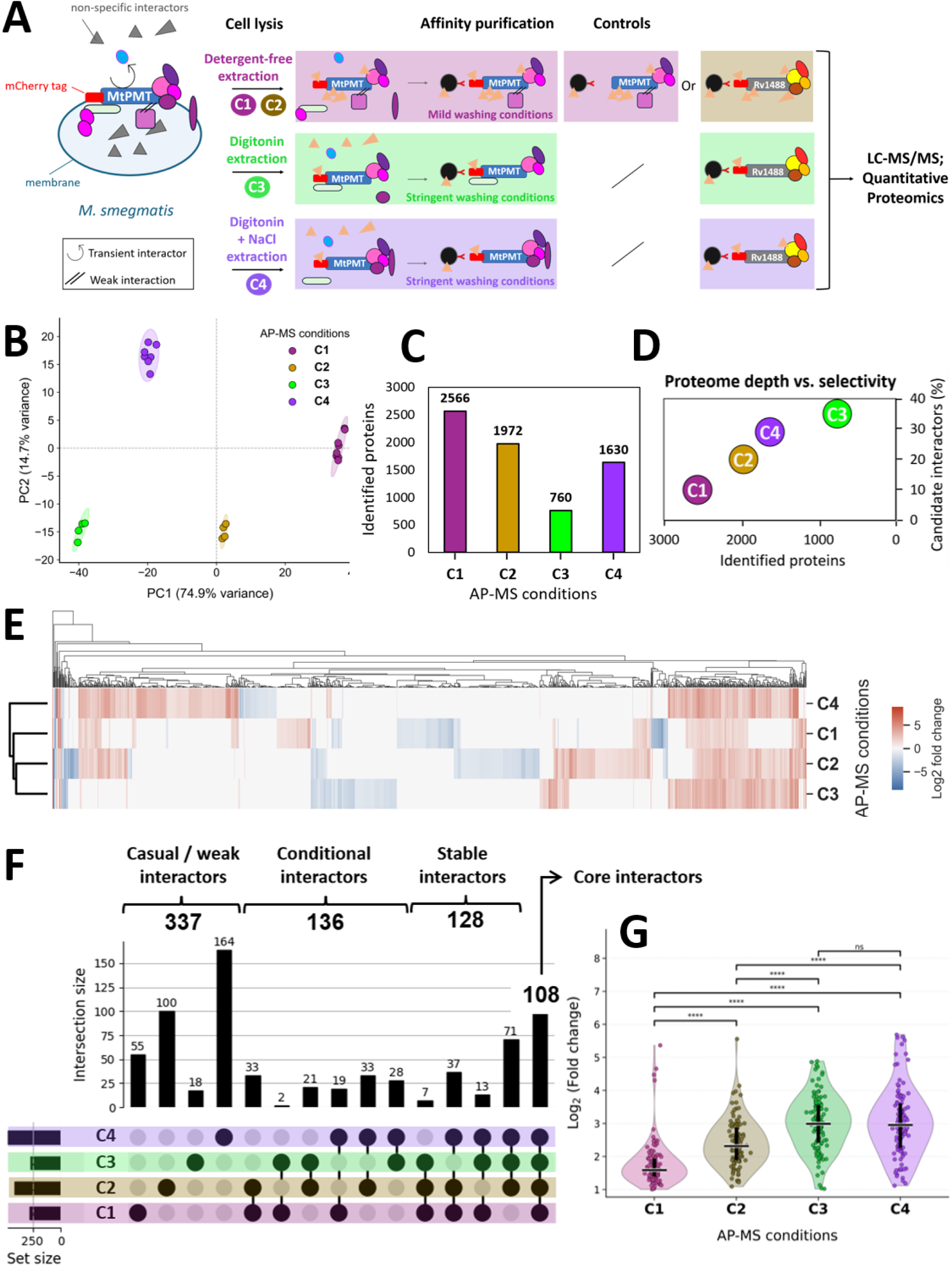
: Biochemical Tuning of AP–MS Reveals Distinct MtPMT-Associated Proteomes but a persistant core. **(A)** Schematic overview of the AP–MS workflow and purification conditions. MtPMT was expressed as an N-terminal mCherry fusion in *M. smegmatis* and subjected to affinity purification (AP) under four conditions of increasing stringency (C1–C4). C1 corresponds to detergent-free lysis under mild conditions of co-immunoprecipitation; C2 incorporates a topology-matched mCherry-tagged membrane protein (Rv1488–mCherry) as specificity control; C3 uses digitonin-mediated membrane solubilization and stringent conditions of co-immunoprecipitation to enrich stable membrane-associated complexes; and C4 combines digitonin extraction with high ionic strength (150 mM NaCl) to further select for highly stable interactions. Eluates were analyzed by LC–MS/MS. Volcano plots corresponding to each condition are shown in **Supplementary Figure 1A**. **(B)** Principal component analysis (PCA) of AP–MS quantitative proteomic profiles across purification conditions (C1–C4). PCA was performed using normalized abundance values from all proteins identified across AP–MS experiments. Replicates clustered tightly within each purification condition, supporting high experimental reproducibility. Separation of conditions along PC1 reflected progressive changes associated with increasing biochemical stringency, whereas PC2 further distinguished digitonin–NaCl extraction (C4) from other purification regimes. Shaded ellipses indicate the 95% confidence interval for each condition. **(C)** Number of proteins identified (either candidate interactors or non-selective proteins) under each AP–MS condition. Mild purification conditions (C1–C2) yield high proteome coverage, whereas digitonin-based extraction (C3–C4) reduces dataset complexity. The proportion of candidate interactors relative to non-selective proteins is shown in **Supplementary Figure 1B**. **(D)** Relationship between interactome depth and selectivity across purification conditions. Increasing stringency results in a progressive shift from high-depth/low-selectivity datasets (C1) toward high-selectivity interactome (C3), with C4 providing an optimal balance between both. **(E)** Hierarchical clustering of protein abundance profiles across AP–MS conditions. Heatmap representation of proteins significantly enriched (|log2FC| ≥ 1, p ≤ 0.05) or depleted (|log2FC| ≤ 1, p ≤ 0.05) across the four AP–MS conditions (C1–C4), based on log2 fold change values. Rows correspond to AP–MS conditions and columns to proteins. Color intensity reflects relative enrichment, with red indicating higher enrichment, blue indicating depletion, and white intermediate values. The row dendrogram clusters conditions according to overall similarity in protein enrichment profiles, revealing a close relationship between C2 and C3, whereas the highest-stringency condition (C4) segregates more distinctly. The column dendrogram groups proteins exhibiting similar enrichment patterns across purification conditions, highlighting subsets of proteins with differential persistence and sensitivity to detergent extraction and elevated ionic strength. Clustering supports a progressive biochemical restructuring of the recovered MtPMT-associated proteomic landscape across purification stringencies. **(F)** Stratification of the MtPMT interactome based on interaction persistence across purification conditions. UpSet plot shows the overlap between datasets, highlighting condition-specific and shared interactors. Proteins were classified into four categories according to their detection across conditions: casual/weak interactors (detected in a single condition), conditional interactors (two conditions), stable interactors (three conditions), and core interactors (all four conditions). The core interactome comprises 108 proteins consistently detected across all conditions, representing the most robust MtPMT-associated partners. **(G)** Global distribution of enrichment scores for the 108 proteins constituting the MtPMT core interactome. Violin plots represent the distribution of log₂(fold change) values obtained under the four AP–MS conditions (C1–C4). Each dot corresponds to an individual protein. The central horizontal black bar indicates the median, and vertical lines represent the interquartile range. A progressive increase in enrichment is observed with increasing purification stringency, reflecting selective retention of robust MtPMT-associated proteins. Statistical comparisons between conditions were performed using paired two-sided Wilcoxon signed-rank tests, followed by Bonferroni correction for multiple comparisons. Significance levels are indicated as ns (not significant); **** P < 0.0001.

In the baseline condition (C1), bacterial cell lysis and mC-MtPMT immunopurification were performed under detergent-free mild conditions designed to preserve the broadest possible range of molecular interactions, including weak and potentially transient associations. Enrichment of mC-MtPMT copurified proteins was assessed by quantitative proteomic relative to an untagged MtPMT specific control expressed by an isogenic strain processed in parallel.

Condition C2 was conducted under similar biochemical conditions but using an alternative control consisting in a similar recombinant *M. smegmatis* expressing an mCherry-tagged membrane protein (Rv1488-mCherry) unrelated to MtPMT. This membrane protein control was introduced to improve discrimination between bona fide MtPMT-associated proteins and nonspecific components arising from the membrane background or affinity purification of mCherry associated proteins.

Conditions C3 and C4 retained the same Rv1488–mCherry reference control but progressively increased purification stringency. In C3, digitonin, a mild non-ionic detergent, was introduced to improve membrane solubilization and facilitate recovery of membrane-associated assemblies. C4 further increased stringency through elevated ionic strength (150 mM NaCl), thereby destabilizing molecular assemblies predominantly maintained by electrostatic interactions.

Principal component analysis (PCA) of quantitative proteomic profiles obtained across C1–C4 confirmed both the reproducibility of the experiments and the progressive restructuring of the MtPMT-associated proteome under distinct biochemical conditions (**Fig. 1B**). Biological replicates clustered tightly within each experimental condition assessing limited intra-condition variability supporting strong experimental robustness. In contrast, proteomic profiles segregated clearly between conditions in multidimensional space. The first principal component represented the major source of proteomic variation (PC1; 74.9% of explained variance) and strongly separated C1 from C3 and, to a lesser extent, from C2 and C4 (**Fig. 1B**). Notably, the separation of low-stringency conditions (C1 and C2) from detergent-based purifications (C3 and C4) along PC1 indicates that detergent-mediated extraction profoundly altered the composition of the recovered MtPMT-associated proteome. The second principal component (PC2; 14.7%) further distinguished C4 from the remaining conditions (**Fig. 1B**). The emergence of a specific proteomic signatures associated with increased purification stringencies indicates that biochemical parameters profoundly shaped the recovered interactant landscape.

Quantitative analysis of the individual proteomic datasets further highlighted the impact of purification conditions on the recovered interactomes (**Fig. 1C–D and Fig. S1**). Condition C1 yielded the deepest proteome, with more than 2,500 proteins identified; however, only 10.7% met both enrichment and statistical significance criteria (log₂ fold change ≥ 1; *p* ≤ 0.05), reflecting limited selectivity. Introduction of the membrane-associated Rv1488–mCherry control in C2 improved discrimination increasing proportion of significantly enriched interactors to 20.8% while maintaining substantial proteome depth (1,972 proteins detected).

The use of digitonin in C3 produced a marked shift toward increased selectivity, with 268 significantly enriched MtPMT candidate partners among the 760 detected proteins (35.3%), albeit at the cost of reduced proteome depth. Finally, combining digitonin with elevated ionic strength in C4 preserved high selectivity while partially restoring proteome depth relative to C3. Collectively, these results demonstrate that purification conditions strongly modulate the propertries of the recovered MtPMT’s interactomes, with digitonin alone (C3) providing maximal selectivity and digitonin combined with increased ionic strength (C4) offering the most balanced compromise between interactome depth and selectivity.

Further comparison of MtPMT associated proteomes identified across the four AP–MS datasets confirmed that purification conditions qualitatively influenced the nature of the recovered partners. Hierarchical clustering of protein abundance profiles revealed distinct sets of proteins displaying differential recovery patterns across purification conditions (**Fig. 1E**), consistent with variable biochemical stability and sensitivity to extraction parameters. Notably, C2 and C3 displayed closely related profiles, indicating that the use of a topology-matched membrane control and digitonin-mediated solubilization provided largely overlapping MtPMT-associated proteomes (**Fig. 1E**). In contrast, C4 segregated more distinctly from the other conditions, consistent with the selective retention of interaction partners resistant to both detergent extraction and elevated ionic strength.

Together, these observations support the existence of a progressive transition in the composition of MtPMT-associated proteomes as purification stringency increases.

To further exploit the complementary information provided by the four AP–MS datasets, we sought to move beyond by classifiying proteins according to their recurrence across experimental conditions. UpSet intersection analysis [45] of the distribution of the 709 proteins fulfilling enrichment and statistical significance selection thresholds across the four AP-MS experiments, evidenced obvious overlaps between the different condition-dependent MtPMT associated proteomes (**Fig. 1F**). Interestingly 108 proteins were found consistently enriched in all four experimental conditions. Because these proteins remained associated with MtPMT even under the most stringent purification conditions, they correspond to the highest-confidence potential partners that define the core MtPMT-associated proteome. Progressive purification stringency resulted in a global shift of the core interactome toward higher enrichment values (log₂[fold change]), confirming that these proteins are preferentially retained under increasingly stringent biochemical conditions (**Fig. 1G**).

Beside these 108 potential core interactors (**Fig. 1F**), 337 proteins were detected exclusively in single experimental condition. These proteins likely should correspond either to peripheral components of the MtPMT molecular environment or to interactions particularly sensitive to biochemical purification parameters—hereafter referred to as casual / weak interactors. Between these two extremes, a substantial fraction of proteins was recovered in either two or three of the four datasets. These were considered conditional interactors and stable interactors, respectively according their intermediate degree of conservation across experimental conditions proteomes.

However, although increasing purification stringency was expected to progressively refine the MtPMT-associated proteome, the different datasets were not strictly nested. Several proteins identified under highly stringent conditions were absent from less stringent datasets, whereas others were selectively recovered only under specific biochemical settings. Such behavior likely reflects both the dynamic nature of membrane-associated assemblies and the intrinsic stochasticity of large-scale proteomic analyses.

### Integrated interactomic unveils MtPMT interaction network architecture driving forces

Rather than neglecting or excluding this inherent inconsistency, we leveraged it to develop a cumulative persistence score (CPS) integrating two complementary dimensions of interaction robustness: (i) the recurrence of protein detection across independent AP–MS experiments and (ii) the relative stringency of the biochemical conditions under which the protein remained associated with MtPMT.

This was achieved by assigning increasing weights ranging from 1 to 4 to the four experimental conditions according to their analytical and biochemical rigor (C1 < C2 < C3 < C4), such that detections under more stringent conditions contributed proportionally more to the final score (**Fig. 2A**). Accordingly, analysis of CPS distribution revealed a continuous spectrum of interaction robustness across the MtPMT-associated proteome (**Fig. 2A**). Core interactors displayed the highest CPS values, consistent with their persistence across all purification conditions, whereas proteins restricted to a single experimental setting occupied the lower end of the distribution. Stable and conditional interactors populated intermediate regions of the scale, indicating progressive differences in biochemical resilience and detection reproducibility.

**Figure 2.**
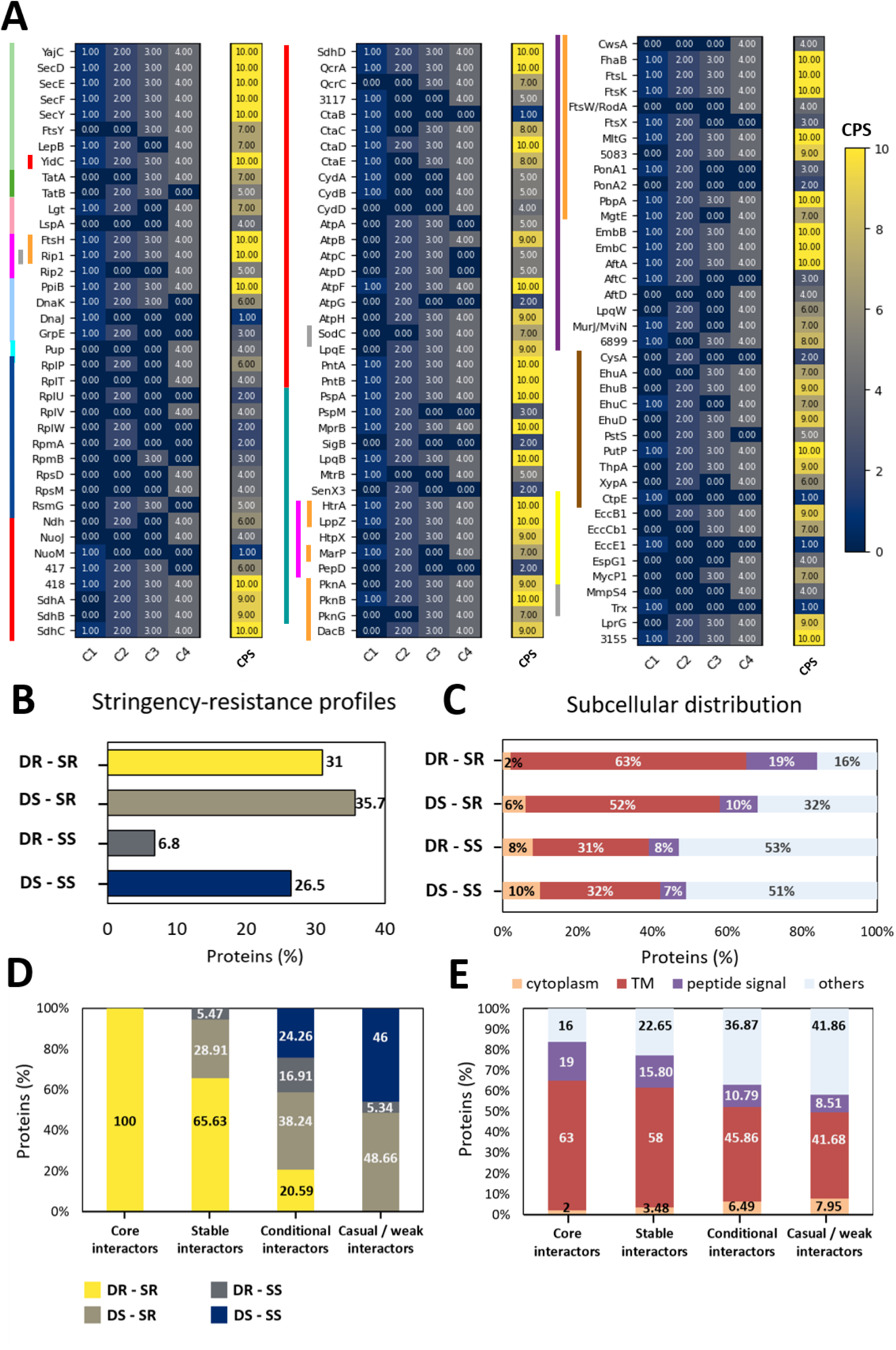
: Integrated interactomic unveils MtPMT interaction network architecture driving forces. **(A)** Quantitative mapping of interaction persistence across AP–MS conditions. Each protein is assigned a cumulative persistence score (CPS) based on its detection across purification conditions of increasing stringency (C1–C4), with higher weights attributed to more stringent conditions. Heatmaps display detection profiles across conditions and corresponding CPS values, revealing a continuous gradient of interaction stability across functional modules of the MtPMT interactome. The heatmap highlights a subset of functionally well-characterized interactors and does not represent the full dataset. Functional categories of the interactors are annotated alongside, and the color code is identical to that used in **Fig. 3** (Sec-dependent translocation in light green, Tat pathway in dark green, lipoprotein maturation in light pink, ribosome in dark blue, membrane proteases in magenta, chaperones in light blue, pupylation pathway components in cyan, two-component regulatory systems in dark bluish green, ABC transporters in brown, cell envelope biogenesis in purple, cell division in orange, ESX-1 secretion system in yellow, respiratory metabolism in red, and cell redox homeostasis in gray). **(B)** Classification of MtPMT-associated proteins according to their resistance to biochemical perturbations. Interactors were grouped into four profiles based on their sensitivity to detergent solubilization and elevated ionic strength: detergent and salt resistant (DR-SR; yellow), detergent sensitive and salt resistant (DS-SR; light green), detergent resistant and salt sensitive (DR-SS; dark green), and detergent and salt sensitive (DS-SS; dark blue). The distribution of proteins across these categories is shown as percentage of total interactors. The criteria used to define resistance profiles are described in **Supplementary Figure 2**. **(C)** Subcellular distribution of partners across resistance profiles. Proteins were classified based on predicted localization features (Uniprot database), including transmembrane helices (TM, red), signal peptides (purple), cytoplasmic localization (orange), and other categories (light blue). Detergent and salt resistant interactions are strongly enriched in membrane proteins, whereas detergent and salt sensitive interactions show increased representation of cytoplasmic proteins. **(D)** Distribution of resistance profiles across interaction strata defined by persistence (core, stable, conditional, and casual interactors). Core interactors are predominantly detergent and salt resistant, whereas casual interactors are enriched in detergent and salt sensitive proteins, indicating a progressive decrease in interaction stability. **(E)** Subcellular distribution across interaction strata. Core and stable interactors are strongly enriched in transmembrane proteins, while conditional and casual interactors display increasing proportions of cytoplasmic and non-membrane-associated proteins. This distribution highlights the coupling between interaction persistence, biochemical resistance, and subcellular localization.

Although this CPS cannot substitute to direct measure of the interaction strength or affinity, it represented a valuable empirical metric reflecting both the reproducibility of protein recovery and its resistance to progressively stronger biochemical perturbations. By integrating information from the four complementary AP–MS experiments this approach converts partially overlapping interactomes into a unified confidence scale, providing an empirical ranking framework for prioritizing MtPMT-associated proteins.

Moreover, CPS ranking allows refining the primary conditions-dependent interactomes by defining four complementary stringency-resistance profiles (**Fig. S2**) namely, “detergent and salt resistant” (DR-SR, 31%), “detergent sensitive and salt resistant” (DS-SR; 35.7%), “detergent resistant and salt sensitive” (DR-SS; 6.8%) and finally “detergent and salt sensitive” (DS-SS; 26.5%) (**Fig. 2B**). Strikingly, this stratification unveiled that majority of MtPMT-associated proteins (66.7%) were resistant to elevated ionic strength indicating that interaction of MtPMT with its associated landscape is predominantly stabilized by hydrophobic or membrane-associated contacts rather than electrostatic interactions.

To further assess whether this stratification reflects biologically meaningful interaction properties, we crossed the stringency-resistance profiles of MtPMT-associated proteins with their structural properties driving their established or predicted subcellular localization. Four protein classes were considered according UniProt annotations [46] (when available) namely: membrane proteins (containing predicted transmembrane helices), secreted or membrane-associated proteins (containing signal peptides), cytoplasmic proteins and undefined. Clear differences emerged across stringency-resistance profiles (**Fig. 2C**). DS-SS interactant subset grossly contained a majority proteins with undefined subcellular localization (51%) together with 10% of cytoplasmic proteins, and 39% of transmembrane and membrane associated proteins. Similar distribution was observed for DR-SS potential partners. In contrast, high ionic strength resistant partner DS-SR subset appeared strongly enriched in membrane-associated proteins. In this fraction, transmembrane proteins represented more than half of all interactors (∼52%), together with an increased representation of proteins containing secretion signal peptides (10%). The strongest enrichment in these latter classes was observed for interactions resistant to both detergent and high ionic strength (DR-SR, subset), where transmembrane and membrane-associated proteins accounted for the vast majority of the interactome (∼82%), whereas cytoplasmic proteins were reduced to less than 2%. Thus, the protein-class composition of these different subsets aligned with their stringency-resistance profile and support the prevalence of the membranous feature of the MtPMT associated interactome.

Similarly, further mapping of the stringency-resistance profiles onto recurrence-defined interactor classes, revealed a strong correlation between interaction robustness, biochemical stability, and subcellular distribution (**Fig. 2D-E**). Core interactors that are exclusively DR-SR proteins, were for their most part either transmembrane proteins or membrane associated proteins, sharing highly stable associations embedded within the membrane environment of MtPMT. Besides, stable interactors displayed a more heterogeneous composition yet dominated by DR-SR components combining membrane-associated proteins with a moderate contribution of less resistant interactions. Finally, conditional and casual/weak interactors showed in contrast, increased representation of DS-SS proteins consistent with a significative enrichment in cytoplasmic partners engaging weak, transient, or indirect interactions with MtPMT. This is exemplified by the fact that cytoplasmic proteins likely interacting with MtPMT through electrostatic associations such as chaperones of the DnaK system (DnaK, DnaJ, GrpE) were detected exclusively under low ionic strength conditions (**Fig. 2A**). Given the central role in protein folding and stress-response proteostasis, the spatial association of these components with MtPMT provides further evidence for a functional coupling between protein O-mannosylation and the cellular protein quality control machinery.

Taken together, these findings indicate that biochemical resistance and interaction persistence are strongly coupled to protein subcellular localization, further underscoring the robustness and discriminatory power of the multi-AP-MS approach. Highly persistent interactions predominantly engage membrane-embedded protein complexes maintained through hydrophobic contacts, whereas lower persistence interaction layers are enriched in cytoplasmic and membrane-associated peripheral proteins. This hierarchical organization supports a model in which MtPMT resides within a stable membrane-embedded core complex, surrounded by progressively more dynamic peripheral interaction networks.

### Functional stratification reveals a membrane-centered multiprocess organization of the MtPMT’s landscape

We then sought to verify whether this organization aligned with a topological functional speciation. However, given that a substantial fraction of MtPMT-associated proteins identified across AP-MS datasets remains poorly or not functionally characterized (**Fig. S3**) our analysis therefore focused only on the well-characterized interactors.

Functional enrichment analysis of proteins identified across all experiments revealed a significant overrepresentation of cellular processes associated with membrane protein biogenesis and cellular metabolism (**Fig. 3A and Fig. S4 A**). Prominent among these were components of the Sec-dependent protein translocation machinery (light green). This finding is consistent with previous observations in *S. cerevisiae*, where PMT enzymes associate with the Sec61 translocon component [24]. In addition to the Sec translocon machinery, multiple components of the twin-arginine translocation (Tat) pathway (dark green) were also detected, in line with recent reports identifying Tat pathway as both transporters of- and targets of PMT-mediated *O*-mannosylation [20] [41] [47].

**Figure 3:**
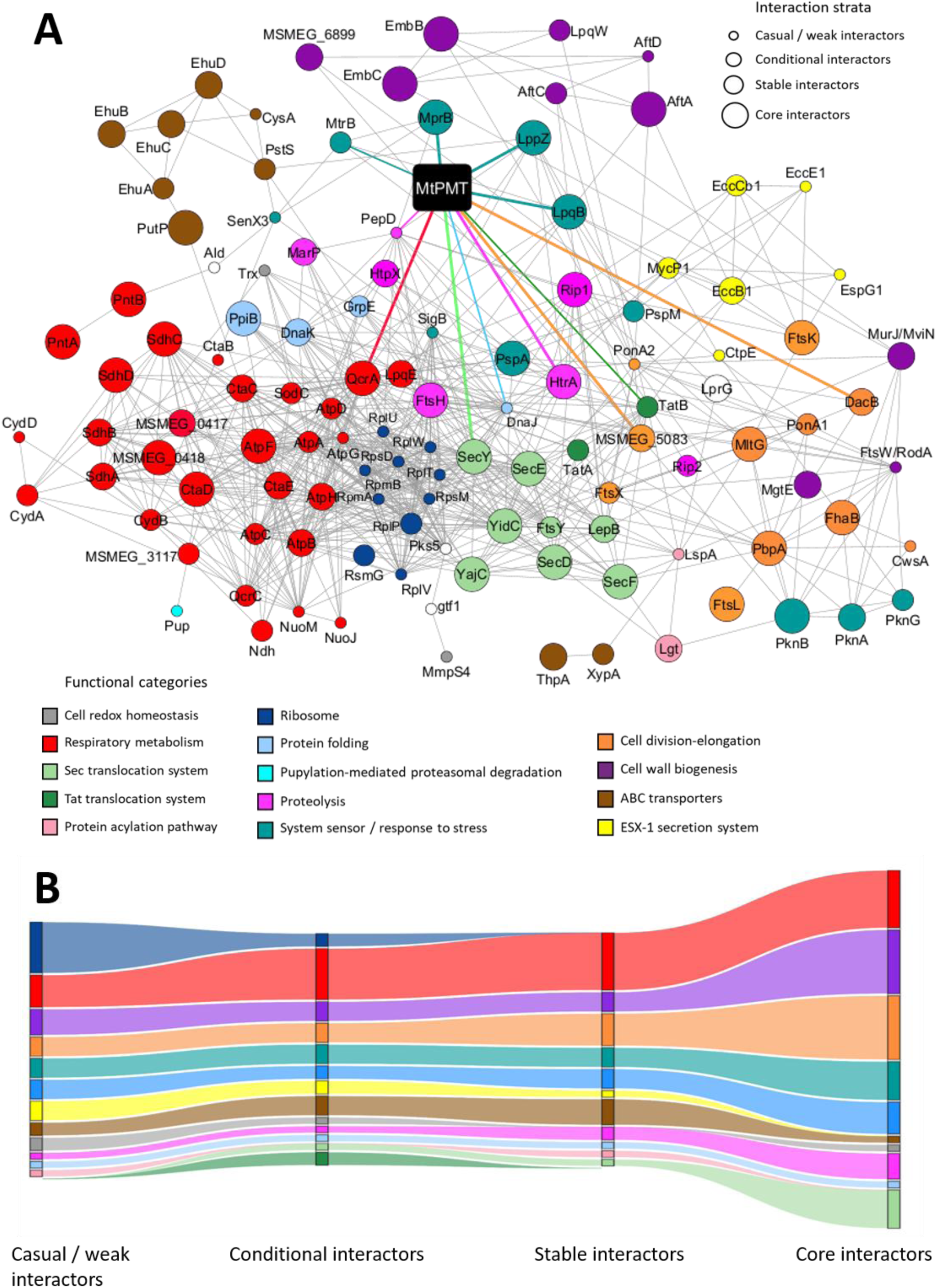
Functional stratification reveals a membrane-centered multiprocess organization of the MtPMT’s landscape. **(A)** Network representation of the MtPMT interactome reconstructed from AP–MS datasets. Nodes represent proteins and edges indicate physical and/or functional associations based on STRING database analysis. Node size is proportional to the persistence of interactor identification, with larger nodes corresponding to core interactome components and smaller nodes to casual or weak interactors. Proteins are color-coded according to functional categories identified by KEGG and Gene Ontology enrichment analyses (**Supplementary Figure 4A**) and by Uniprot and Mycobrowser annotations: Sec-dependent translocation (light green), Tat pathway (dark green), lipoprotein maturation (light pink), ribosome (dark blue), proteostasis factors including membrane proteases (magenta), chaperones (light blue), and pupylation pathway components (cyan), as well as two-component regulatory systems (dark bluish green), ABC transporters (brown), cell envelope biogenesis (purple), cell division (orange), ESX-1 secretion system (yellow), respiratory metabolism (red), and cell redox homeostasis (gray). Colored edges indicate predicted functional associations between MtPMT and its identified partners based on STRING database analysis. **(B)** Functional stratification of MtPMT-associated proteins across interaction classes. Sankey diagrams show the distribution of enriched functional categories among four interaction layers: casual/weak interactors, conditional interactors, stable interactors, and core interactome. The detailed list of proteins assigned to each functional category and interaction class is provided in **Supplementary Figure 5.**

Proteins involved in lipoprotein maturation, (e.g., the diacylglyceryl transferase Lgt and the signal peptidase LspA) were also enriched (**Fig. 3A**; light pink), further linking MtPMT to the post translational lipidation of secreted bacterial manno-lipoproteines. Furthermore, KEGG pathway analysis [48] highlighted enrichment in ribosomal components (**Fig. 3A and Fig. S4 A**; dark blue), supporting the integration of MtPMT in a molecular environment coordinated with protein synthesis, post-modifications and membrane targeting.

These findings strongly support the current model in which the mannosylation process takes place in the close vicinity of the mycobacterial protein elongation/secretion and lipidation system [40].

Moreover, beyond these predicted association with protein translocation/post-translational modification machinery, the MtPMT interactome was surprisingly found strongly enriched in components of membrane proteostasis and quality control systems (**Fig. 3A**). These included two-component regulatory system factor (e.g., MprB; dark bluish green), membrane-associated proteases (FtsH, Rip1, HtrA, HtpX, MarP, PepD; pink), molecular chaperones (DnaK, DnaJ, GrpE; light blue), and components of the pupylation pathway (Pup; cyan). The enrichment of these functionally diverse yet mechanistically interconnected quality control components, provides compelling evidence that MtPMT is physically embedded within the mycobacterial membrane proteostasis network, suggesting that O-mannosylation may mechanistically contributes to the surveillance, maintenance, and turnover of membrane protein complexes.

Additional enriched functional categories included proteins involved in ABC transport (brown), cell envelope biogenesis (purple), and cell division processes (orange) (**Fig. 3A**), indicating a broader integration within membrane-associated cellular functions. Notably, components of the ESX-1 secretion system were also detected (yellow). Given that several *Mtb*’s ESX-associated proteins have been previously identified as mannoproteins [20] [41], this suggests a potential link between the *Mtb*’s virulence associated Type VII secretion system and protein O-mannosylation.

Strikingly, KEGG [48] and Gene Ontology [49] enrichment analyses revealed also a substantial contribution of proteins involved in respiratory metabolism, including components of the electron transport chain and oxidative phosphorylation (**Fig. 3A and Fig. S4 A**; red). This unexpected enrichment points out a potential spatial clustering of MtPMT with membrane associated respiratory complexes, raising the possibility that O-mannosylation may influence respiratory function and redox homeostasis.

Deeper examination of these cellular process’s distribution across recurrence-defined interactor classes further revealed a gradual functional reorganization along the interaction-strength gradient, underscoring the marked functional stratification of the MtPMT interactome (**Fig. 3B and Fig. S5**).

At the periphery of the interaction landscape, the casual and weak interactors were associated with a broad and heterogeneous set of functions. These included ribosomal proteins, redox homeostasis factors, and proteins linked to general cellular processes, consistent with either low-affinity interactions or indirect long-range interactions. Notably, respiration and protein secretion pathways were only weakly represented in this layer.

In contrast, the conditional interactors displayed a marked enrichment in transport and secretion-associated functions, including ABC transporters and Tat and ESX-1 secretion system components. This layer also showed increased representation of stress-response and envelope-associated processes, figuring a transition from non-specific associations toward functionally relevant membrane processes.

Stable interactors further refined this organization, with a stronger enrichment in core cellular functions associated with membrane physiology. These included cell division and elongation (MSMEG_5083, PonA1, DacB), cell envelope biogenesis, and proteostasis-related pathways, alongside an increased contribution of respiratory metabolism. Compared to conditional interactors, this layer exhibited a more focused functional profile, consistent with increased interaction stability.

The core interactome represented the most functionally specialized layer in which proteins involved in membrane protein biogenesis (Sec system), proteostasis (MprB, HtrA-LppZ, FtsH, Rip1), respiratory metabolism, and cell division were most highly enriched.

Collectively, these results confirm that MtPMT is far from being an isolated enzyme but rather fully integrated within a highly organized membrane-centered, multiprocess environment. General cellular processes and cytoplasmic functions were predominantly confined to peripheral interactions, whereas membrane-associated pathways—including secretion, envelope biogenesis, proteostasis, and respiration—became progressively enriched with increasing interaction stability. This hierarchical organization indicates that interaction persistence is coupled to functional specialization within the MtPMT network.

### MtPMT’s interactome undergoes growth stage-dependent spatio-temporal remodeling

We then sought to verify whether this platform undergoes dynamic reorganization during bacterial growth. To this end, we compared the integrated MtPMT interactome reconstructed from the four AP–MS conditions at late stationary phase (96 h) with an independent interactomic dataset obtained during an earlier physiological stage (72 h), corresponding to early stationary phase. Importantly, the 72 h analysis was performed under biochemical conditions equivalent to the most stringent purification regime (C4), including membrane controls, digitonin solubilization, and elevated ionic strength, thereby enriching for robust associated interactions. This design allowed us to assess how the composition of the MtPMT-associated platform evolves across growth stages while minimizing the contribution of weak interactions.

Comparative analysis revealed a substantial remodeling of the MtPMT interaction landscape between these two stages (**Fig 4A, Table 1 and Fig. S4 B**). While a core set of interactors remained conserved, including key components of the Sec system (SecD, SecE, SecF, SecY, YajC, FtsY, YidC) and the membrane protease FtsH, multiple functional categories displayed marked stage-dependent enrichment, indicating a dynamic reorganization of the MtPMT-associated platform (**Table 1**).

**Figure 4:**
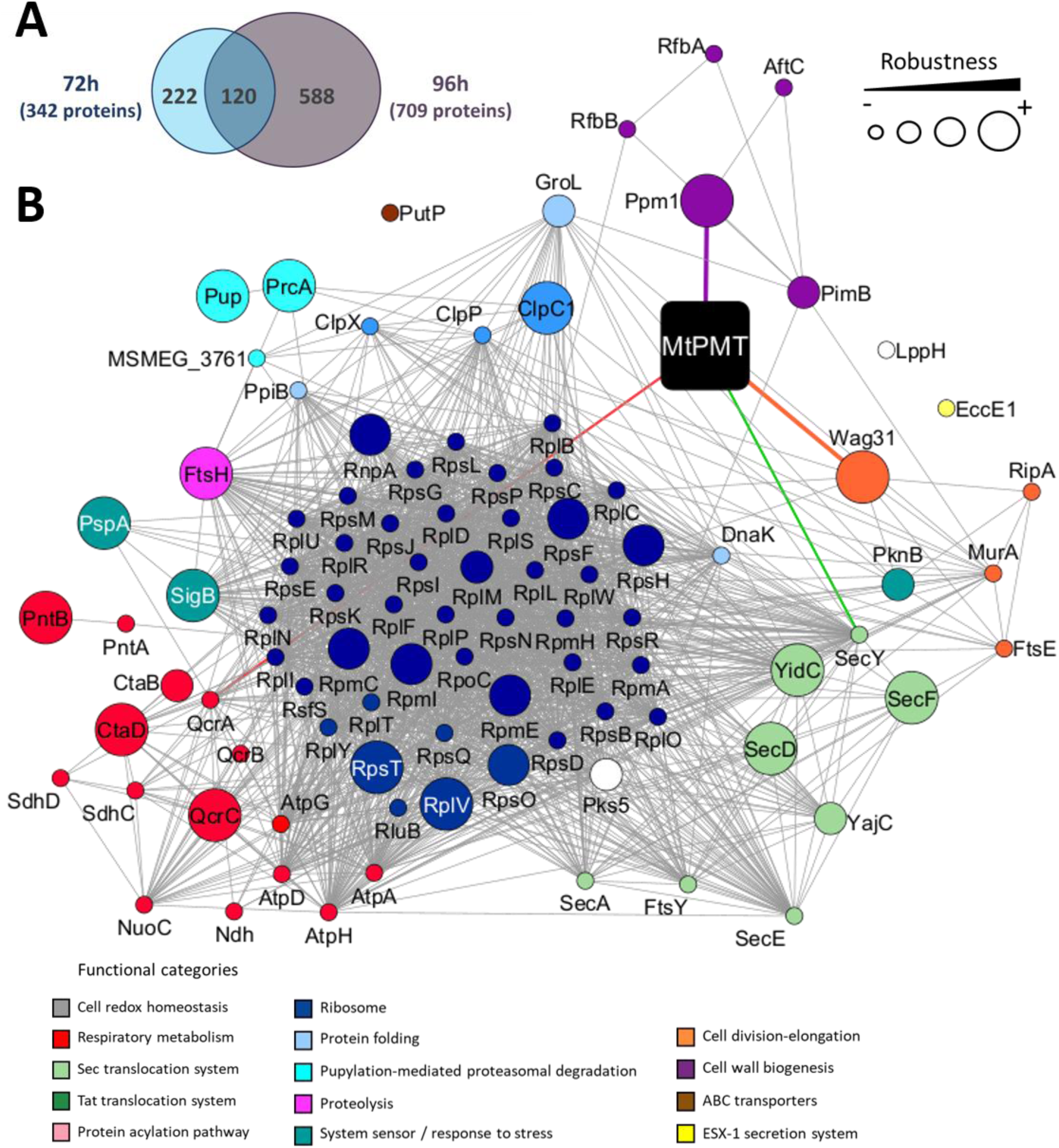
MtPMT’s interactome undergoes growth stage-dependent spatio-temporal remodeling. **(A)** Venn diagram showing the overlap between MtPMT-associated proteins identified by AP–MS at early (72 h) and late (96 h) stationary phases. The number of shared and stage-dependent interactors is indicated, highlighting partial conservation of the interactome across growth stages, together with the total number of proteins identified at each stage annotated next to the diagram. For the 72 h condition, two independent successive experiments were performed, and interactors from both experiments were combined, resulting in a total of 342 proteins. For the 96 h condition, interactors identified across the four independent AP–MS analyses (C1–C4) were combined, yielding a total of 709 proteins. A partial list of functionally characterized proteins belonging to shared and stage-specific fractions is provided in **Table 1**. **(B)** Network representation of the MtPMT interactome at 72 h. Nodes represent proteins and edges indicate predicted physical and/or functional associations (STRING annotations). Colored edges indicate predicted functional associations between MtPMT and its identified partners. Node size reflects the robustness of interactor detection across experiments and enrichment levels: large nodes correspond to proteins identified in both experiments; intermediate nodes correspond to proteins identified in a single experiment with high enrichment (log₂ fold change ≥ 4); smaller nodes correspond to proteins identified in a single experiment with moderate enrichment (log₂ fold change ≥ 2); and the smallest nodes correspond to proteins identified in a single experiment with log₂ fold change between 1 and 2. Proteins are color-coded identically to that used in **Fig. 3**, according to functional categories identified by KEGG and Gene Ontology enrichment analyses (**Supplementary Figure 4B**) and by Uniprot and Mycobrowser annotations. The network is characterized by a strong enrichment in ribosomal proteins and factors associated with protein synthesis, secretion, and proteostasis, illustrating the organization of the MtPMT-associated platform at this stage.

**Tableau 1.**
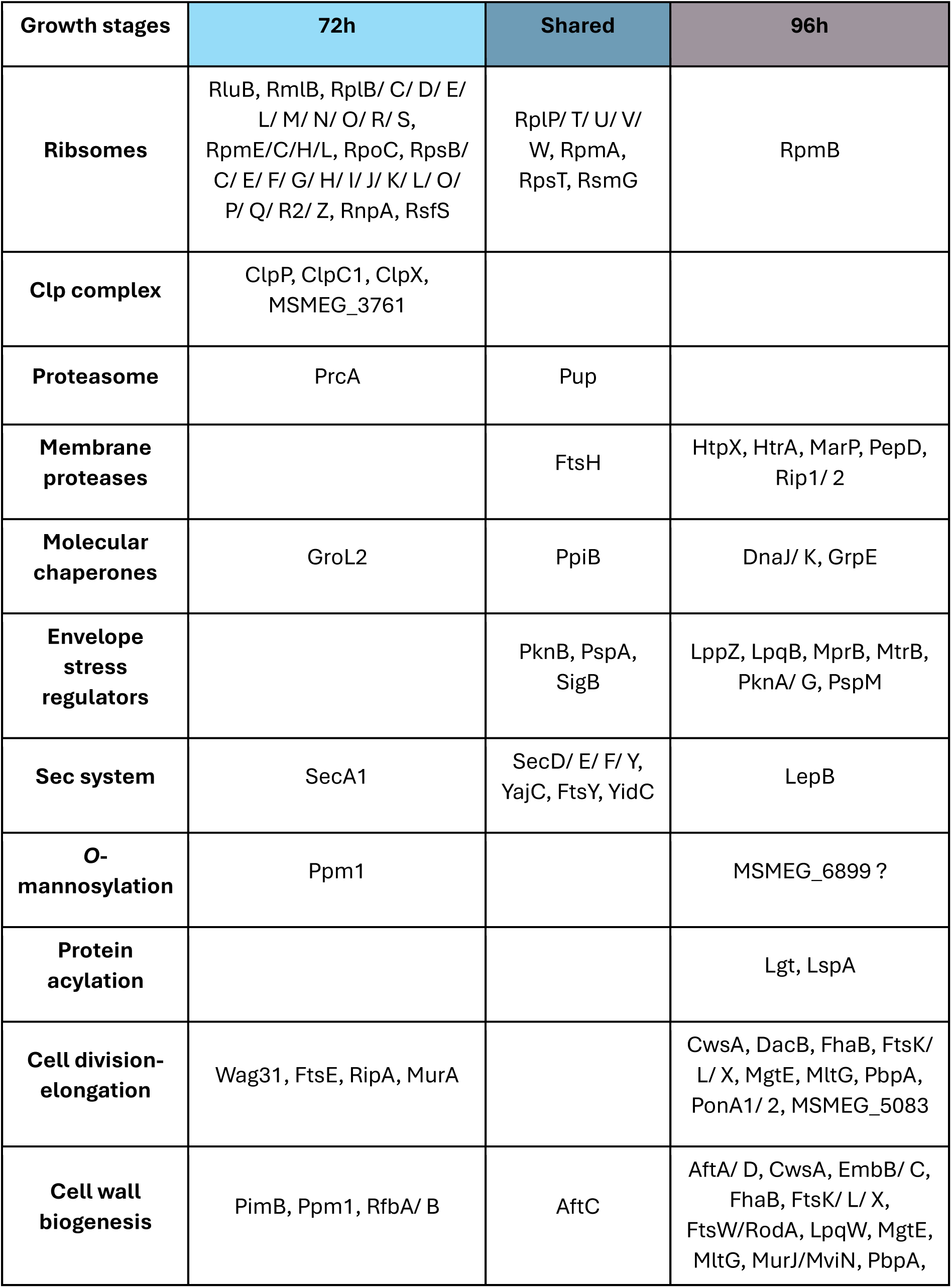
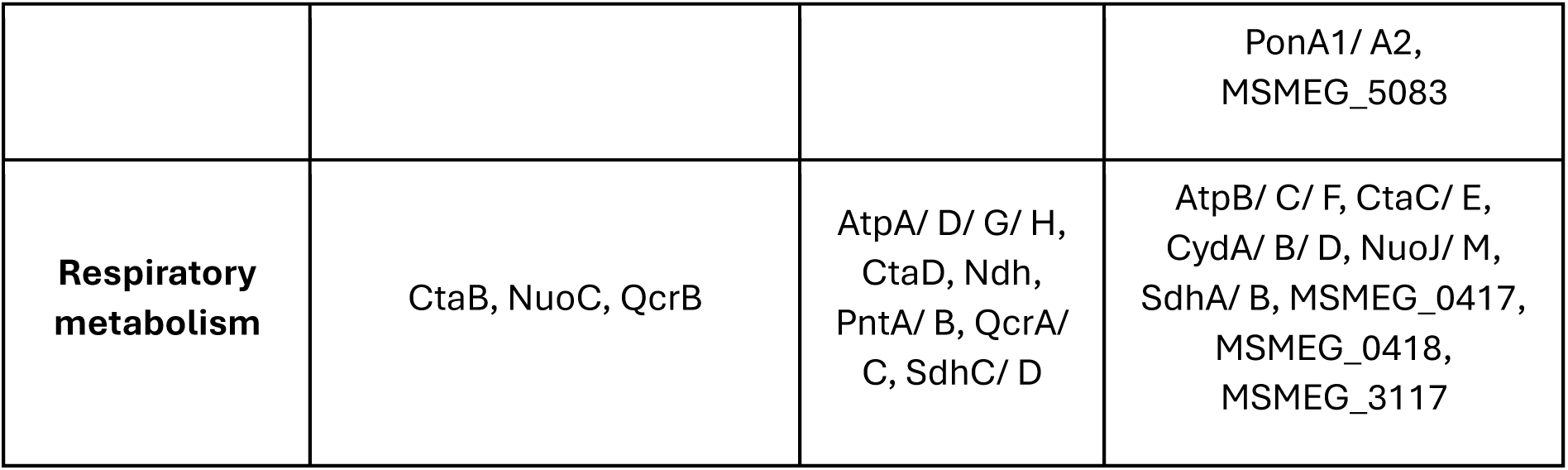
: Functional classification of shared and growth stage-dependent MtPMT interactors. MtPMT-associated proteins identified at 72 h and 96 h, grouped by functional category and growth stages (stage-specific for 72 h or 96h, and shared interactors between both). This list corresponds to the Venn diagram shown in **Figure 4A**.

| Growth stages | 72h | Shared | 96h |
| --- | --- | --- | --- |
| **Ribosomes** | RluB, RmlB, RplB/ C/ D/ E/ L/ M/ N/ O/ R/ S, RpmE/C/H/L, RpoC, RpsB/ C/ E/ F/ G/ H/ I/ J/ K/ L/ O/ P/ Q/ R2/ Z, RnpA, RsfS | RplP/ T/ U/ V/ W, RpmA, RpsT, RsmG | RpmB |
| **Clp complex** | ClpP, ClpC1, ClpX, MSMEG_3761 |  |  |
| **Proteasome** | PrcA | Pup |  |
| **Membrane proteases** |  | FtsH | HtpX, HtrA, MarP, PepD, Rip1/ 2 |
| **Molecular chaperones** | GroL2 | PpiB | DnaJ/ K, GrpE |
| **Envelope stress regulators** |  | PknB, PspA, SigB | LppZ, LpqB, MprB, MtrB, PknA/ G, PspM |
| **Sec system** | SecA1 | SecD/ E/ F/ Y, YajC, FtsY, YidC | LepB |
| **O-mannosylation** | Ppm1 |  | MSMEG_6899 ? |
| **Protein acylation** |  |  | Lgt, LspA |
| **Cell division-elongation** | Wag31, FtsE, RipA, MurA |  | CwsA, DacB, FhaB, FtsK/ L/ X, MgtE, MltG, PbpA, PonA1/ 2, MSMEG_5083 |
| **Cell wall biogenesis** | PimB, Ppm1, RfbA/ B | AftC | AftA/ D, CwsA, EmbB/ C, FhaB, FtsK/ L/ X, FtsW/RodA, LpqW, MgtE, MltG, MurJ/MviN, PbpA, |
|  |  |  | PonA1/ A2,<br>MSMEG_5083 |
| **Respiratory**<br>**metabolism** | CtaB, NuoC, QcrB | AtpA/ D/ G/ H,<br>CtaD, Ndh,<br>PntA/ B, QcrA/<br>C, SdhC/ D | AtpB/ C/ F, CtaC/ E,<br>CydA/ B/ D, NuoJ/ M,<br>SdhA/ B, MSMEG_0417,<br>MSMEG_0418,<br>MSMEG_3117 |

At 72h, the MtPMT interactome was predominantly enriched in factors associated with active protein synthesis and translocation (**Fig 4B, Table 1 and Fig. S4 B**). Notably, SecA1—the cytoplasmic ATPase driving Sec-dependent translocation—was specifically detected at this stage, together with Ppm1, the enzyme responsible for synthesizing the lipid-linked mannose donor required for O-mannosylation. In addition, ribosomal proteins and translation-associated factors were much more abundant, indicative of a spatial coupling between protein-synthesis, - targeting, and -translocation. The concomitant detection of ribosomal proteins, SecA1, translocon components, and Ppm1 co-purified with MtPMT indicates that protein synthesis, secretion, and glycosylation must be coordinated within a shared membrane microenvironment. Consistently, proteolytic systems, including the Clp protease complex (ClpP, ClpX, ClpC1) and the proteasome (PrcA), were also enriched at this stage, indicating a protein quality control probably associated with mannosylation contributing to proteostasis during sustained metabolic activity.

In contrast, at 96 h, the interactome displayed a pronounced shift toward protein maturation and stress adaptation processes (**Table 1**). While core components of the Sec system remained associated, there was an increased representation of proteins involved in post-translocational processing, including lipoprotein maturation (Lgt, LspA) and signal peptide cleavage or stress-responsive proteolysis (LepB, Rip1, HtrA-LppZ). Molecular chaperones (DnaK, DnaJ, GrpE) and regulators of envelope stress responses (MprB, PknB) were also more prominently detected, indicating a membrane environment increasingly dedicated to protein folding, stabilization, and stress management during late stationary phase.

This dynamic reorganization suggests that MtPMT may operate within a flexible membrane platform that adapts to physiological demands, coordinating protein secretion, quality control, and envelope homeostasis according growth phases.

### FtsH and YajC but not MprB are direct spatial neighbors of MtPMT within the membrane platform

Due to the large number of candidate interaction partners identified, even within the restricted core interactome, the individual validation of each physical and functional interaction with MtPMT would represent a considerable undertaking beyond the scope of this study. To assess the reliability of our approach, we therefore selected a subset of proteins for validation based on several criteria: (i) the availability of functional annotation; (ii) to be part of the core interactome; (iii) high enrichment scores across the four AP–MS experiments, particularly in the most stringent one (C4); (iv) a potential functional or genomic association with MtPMT suggested by the literature [50] (**Fig. 3A**); (v) representation of distinct physiological pathways or cellular modules; and (vi) availability of genetic resources, particularly through the Mycobacterial Systems Resource Database (MSRdb) [51].Based on these criteria, we first focused on three candidate partners, YajC, FtsH, and MprB, which are among the best-characterized proteins displaying the highest average enrichment scores (**Fig. 5A and Fig. S6**). Their strong and consistent enrichment across all experimental conditions, including under the stringent C4 condition (**Fig. 5A**), further supports their robust association with MtPMT. Importantly, these proteins participate in distinct physiological processes—protein secretion and membrane biogenesis (YajC), membrane proteostasis (FtsH), and envelope stress response (MprB)—thereby providing an opportunity to explore the integration of MtPMT within diverse membrane-associated cellular networks. As recombinant Dendra2–Flag fusion constructs for these proteins were available through the MSRdb [51], we first sought to validate their physical association with MtPMT by reciprocal affinity purification. To this end, Dendra2–Flag-tagged versions of YajC, FtsH, and MprB were independently co-expressed with mC–MtPMT in *M. smegmatis* and immunopurified using anti-Flag affinity resin. As shown in **Fig. 5B**, mC–MtPMT was specifically detected in the eluates using anti-mCherry nanobodies. The selective reciprocal co-purification of mC–MtPMT with YajC, FtsH, and MprB thus confirms these proteins as bona fide interaction partners and prominent components of the MtPMT membrane-associated interactome.

**Figure 5:**
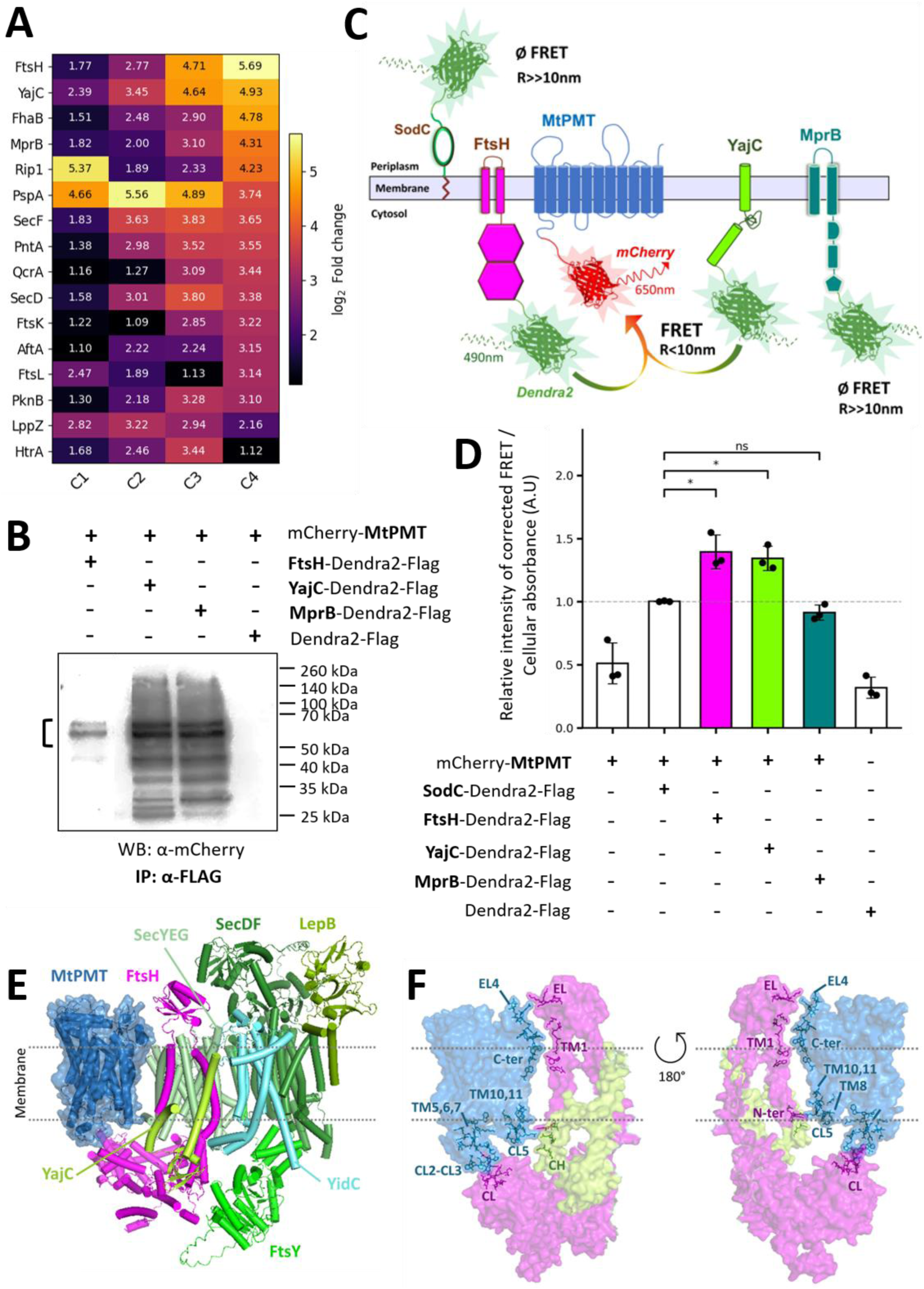
FtsH and YajC but not MprB are direct spatial neighbors of MtPMT within the membrane platform. **(A)** Heatmap showing the log₂(fold change) values of selected proteins from the core interactome across the four AP–MS conditions. YajC and FtsH show the highest fold changes under the more stringent condition (C4) together with a consistent stepwise increase across conditions. The complete heatmap of all 108 core interactors is provided in **Supplementary Figure 6**. **(B)** Validation of selected MtPMT interactions by reciprocal affinity purification. Dendra2–Flag fusion constructs of candidate interactors (FtsH, YajC, and MprB) were co-expressed with mCherry–MtPMT in *M. smegmatis* and subjected to anti-Flag immunopurification (IP). Western blot (WB) analysis shows specific co-purification of mCherry– MtPMT in eluates, confirming these proteins as bona fide MtPMT partners. The mCherry–MtPMT fusion protein is detected at the expected molecular weight (50–70 kDa; annotated by parenthesis), while additional bands likely correspond to aggregated or truncated forms of the protein. Expression of the fusion proteins in cell lysates was verified prior to IP by fluorescence measurements in live bacteria or by WB (**Supplementary Figure 7**). “+” and “–” indicate the presence or absence of the indicated constructs, respectively. **(C)** Schematic representation of the FRET experimental design. mCherry fused to MtPMT serves as the acceptor fluorophore, while Dendra2 fused to candidate interactors (FtsH, YajC, MprB) or controls is used as the donor. The periplasmic protein SodC–Dendra2–Flag serves as a negative control due to spatial separation from cytoplasmically oriented mCherry. Energy transfer occurs only when donor and acceptor fluorophores are within ∼10 nm radius (R), enabling detection of spatial proximity *in vivo*. “Ø” indicates absence of detectable FRET signal. **(D)** Quantification of FRET efficiency in live bacteria. Significant FRET signals are observed for FtsH– and YajC–Dendra2–Flag when co-expressed with mCherry–MtPMT, indicating close spatial proximity. In contrast, MprB–Dendra2–Flag and control constructs display background-level signals. Data are presented as corrected FRET intensity normalized to cellular absorbance (arbitrary units, A.U.) and expressed relative to the SodC-Dendra2-Flag control signal. Expression of the fusion proteins in cell lysates was verified prior to FRET analysis by fluorescence measurements in *live* bacteria or by WB (**Supplementary Figure 7**). Statistical comparisons were performed using unpaired two-sided Student’s *t*-tests with Welch’s correction. Significance levels are indicated as ns, not significant; * P < 0.05. **(E)** AlphaFold3 predictive model of the MtPMT within a Sec-associated assembly including YajC, FtsH, and additional Sec components. Proteins are represented in cartoon, with MtPMT additionally shown in surface representation. The membrane is schematically indicated by dashed lines. Alternative views of the global model are provided in **Supplementary Figure 8A**. ipTM = 0.36; pTM = 0.41 **(F)** Structural focus on the MtPMT–FtsH–YajC interfaces. Proteins are displayed as semi-transparent surfaces. Amino acids located within a 10 Å distance between MtPMT and its partners are highlighted, defining putative interaction interfaces. The corresponding structural elements: transmembrane helices (TM), cytoplasmic loops (CL), cytoplasmic helix (CH), and extracytoplasmic loops (EL), involved in these interfaces are annotated. Two orientations (180° rotation) are shown to visualize the spatial organization of the interaction surfaces. An alternative representation of these interfaces is shown in **Supplementary Figure 8B**, and the list of amino acids involved is provided in **Supplementary Table 2**.

To further confirm experimentally these spatial proximities *in vivo*, we performed fluorescence resonance energy transfer (FRET) analyses in live bacteria [52]. Using Dendra2 and mCherry as the fluorescence donor and acceptor respectively, we detected significant FRET signals for recombinant bacteria co-expressing the FtsH– or YajC–Dendra2–Flag constructs with mC–MtPMT (**Fig. 5C-D**). In contrast, the FRET signal observed with MprB–Dendra2–Flag was indistinguishable from the background detected for the negative controls including the periplasmic SodC– Dendra2–Flag or the individual Dendra2–Flag or mCherry expressed intracellularly (**Fig. 5C-D**). These results indicate that the fluorescent tags on the FtsH– or YajC fusion proteins are located within ≤10 nm of the mCherry moiety of mC-MtPMT, consistent with direct physical interaction or very close spatial association *in vivo*.

Thus these findings, consistent with the affinity purification data provide concrete evidences that, in the bacterial cell, FtsH and YajC—which exhibit the highest C4 enrichment (**Fig. 5A**)—are direct spatial neighbors of MtPMT whereas MprB—which shows lower C4 enrichment (**Fig. 5A**)—likely engages more indirectly.

To gain structural insight into the nature of these interactions, we next generated a predictive model of the spatial organization of MtPMT together with its validated partners using AlphaFold3 [53]. Because YajC and the membrane protease FtsH are both functionally linked to Sec-dependent protein translocation and quality-control pathways [54] [55], representative components of the *Mtb* Sec machinery were included in the structural prediction query. Interestingly, the resulting model positioned MtPMT within the Sec-associated complex in close proximity to YajC and to FtsH (**Fig. 5E-F and Fig. S8**).

To further characterize the predicted assembly model, inter-protein distances were analyzed with a focus on contacts within 10 Å, a threshold indicative of potential interaction interfaces. Only one interface below this distance threshold was detected between MtPMT and YajC, connecting the N-terminal extremity of transmembrane helix (TM) 11 of MtPMT with the cytoplasmic helix (CH) of YajC (**Fig. 5F, Fig. S8 B and Table S2)**. By contrast, the assembly model unveiled four distinct interaction interfaces between MtPMT and FtsH: (i) a first interface linking cytoplasmic loop (CL) 5, the C-terminal extremity of TM10, and TM8 of MtPMT to the N-terminal region of FtsH; (ii) a second interface connecting the C-terminal extracytoplasmic region of MtPMT with the C-terminal end of TM1 of FtsH; (iii) a third interface between the extracytoplasmic loop (EL) 5 of MtPMT and the extracytoplasmic loop of FtsH; and (iv) a fourth interface bringing multiple cytoplasmic elements of MtPMT—including CL2 and CL3, the N-terminal region of TM5, the C-terminal region of TM6, and the N-terminal region of TM7—into close proximity with the cytoplasmic domain of FtsH (**Fig. 5F, Fig. S8 B and Table S2).**

Together, these structural predictions are consistent with the affinity purification data and support a model in which MtPMT is integrated into a Sec-associated membrane platform. Within this assembly, YajC and FtsH appear to constitute direct spatial neighbors of MtPMT, whereas the interaction with MprB may be mediated indirectly through the broader membrane-associated protein network.

### Spatial clustering of MtPMT interactors underlies functional integration of distinct cellular pathways

The identification of MtPMT-associated proteins by AP-MS revealed a marked enrichment of factors involved in proteostasis, maintenance of cell envelope integrity, and respiratory chain function. This organization suggested that MtPMT is embedded within a broader membrane-associated network integrating multiple envelope-related processes. These observations raised the possibility that protein O-mannosylation is not an isolated enzymatic activity but instead functionally integrated into distinct physiological pathways through spatial clustering of its associated partners within a membrane platform.

To test this hypothesis, we first assessed whether disruption of selected MtPMT-associated proteins impacts O-mannosylation activity. MtPMT activity was measured *in cellulo* in available ΔftsH, ΔyajC, and ΔmprB mutants by quantifying O-mannosylation levels of the reporter protein FasC using a previously established lectin-based assay [56]. In addition to these validated interactors, we also selected several proteins functionally linked to MtPMT and either experimentally documented or predicted partners (**Fig. 3A**), including HtrA, LppZ [57], and MSMEG_5083, which displayed some of the highest enrichment scores under stringent AP-MS conditions (**Table S1**). These proteins represent distinct physiological modules, including envelope stress response, membrane proteostasis, and cell envelope organization, and are non-essential under standard laboratory conditions, allowing genetic inactivation [51].

Deletion of the direct MtPMT-associated partners ftsH and yajC resulted in a substantial decrease in O-mannosylation activity (∼70–75% of wild-type levels) (**Fig. 6A**). Given that FtsH is a membrane-anchored AAA+ protease involved in protein quality control [55], and YajC is associated with the Sec translocon [54] [58], these results indicate that MtPMT activity depends on its spatial and functional coupling to protein translocation and membrane proteostasis systems. One possible explanation is that FtsH contributes to maintaining the stability or turnover of MtPMT and/or its substrates, while YajC may facilitate the efficient delivery of nascent polypeptides to the glycosylation machinery.

**Figure 6:**
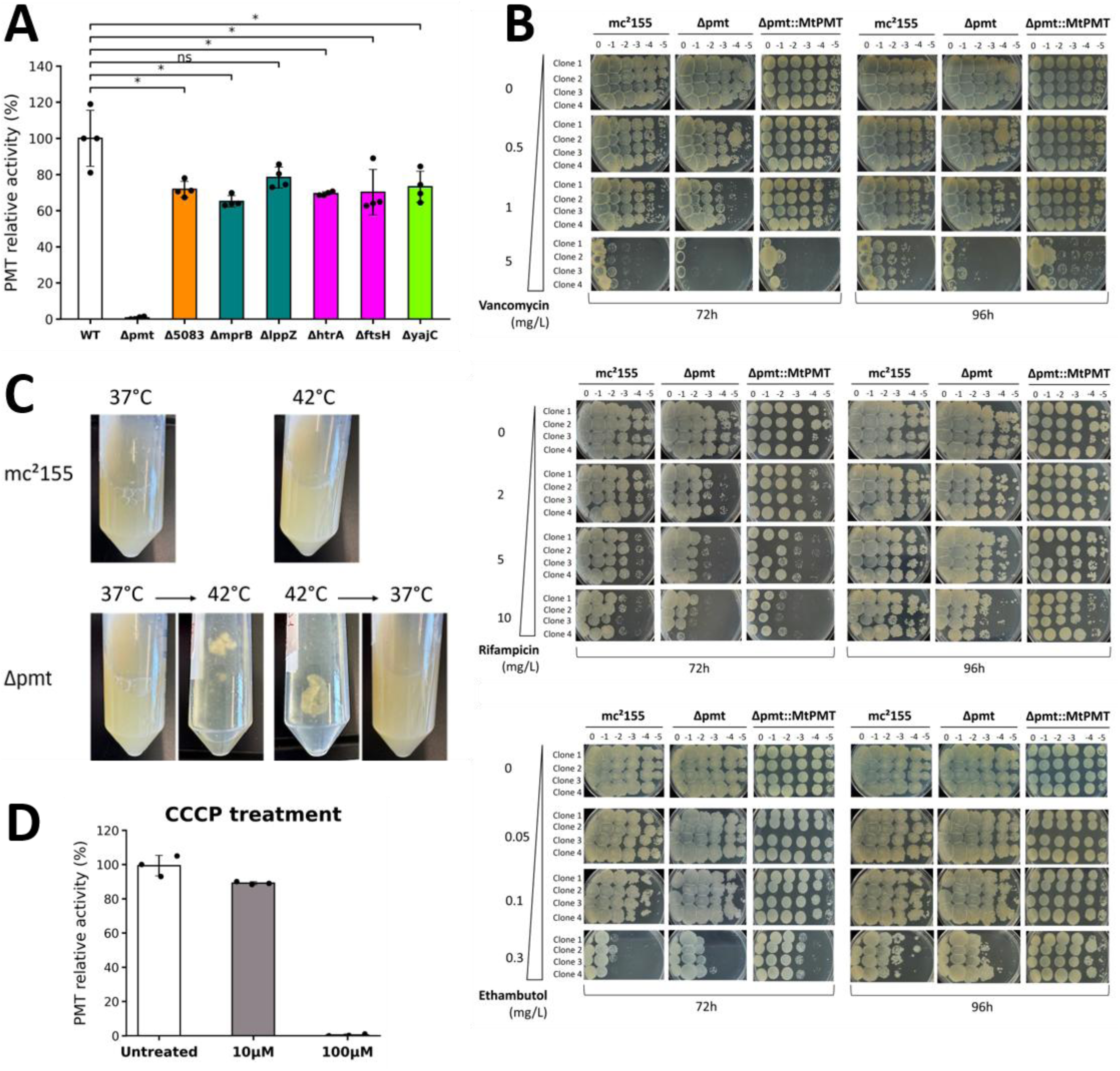
Spatial clustering of MtPMT interactors underlies functional integration of distinct cellular pathways. **(A)** Quantification of PMT activity in *M. smegmatis* knockout strains. *O*-mannosylation activity of the native PMT (MSMEG_5447) was assessed using a lectin-based assay monitoring the glycosylation status of the reporter protein FasC *[1]*. Activity is expressed relative to the wild-type strain (WT). The *Δpmt* strain serves as a negative control, as expected, shows no detectable activity. Deletion of direct spatial neighbors (*ΔftsH*, *ΔyajC*) and selected functionally associated partners (*ΔmprB*, *ΔhtrA*, *ΔlppZ*, *Δmsmeg_5083*) results in variable reductions of PMT activity, supporting their contribution to the MtPMT-associated network. Bars represent mean values ± s.d., and dots indicate four independent biological replicates. Statistical comparisons were performed against the WT strain using unpaired two-sided Welch’s t-tests. Significance levels are indicated as ns, not significant; * P < 0.05. The corresponding FasC quantification is shown in **Supplementary Figure 9**. **(B)** Sensitivity to antibiotics. Serial dilution spotting assays were performed at 72 h and 96 h using WT (*mc²155*), *Δpmt*, and complemented strains (*Δpmt::MtPMT*) in the presence of increasing concentrations of vancomycin, rifampicin, or ethambutol. For each condition, four independent biological clones were analyzed. The *Δpmt* shows increased sensitivity to vancomycin and rifampicin, whereas no significant difference is observed for ethambutol, indicating a selective alteration of envelope permeability to bulky antibiotics. Complementation restores wild-type resistance levels. **(C)** Growth phenotype under thermal stress. Cultures of mc²155 and *Δpmt* strains were incubated at 37°C or 42°C. Representative images from four independent biological clones are shown. The *Δpmt* mutant exhibits a pronounced aggregation phenotype at 42°C. This phenotype is reversible and temperature-dependent, consistent with altered cell surface properties under stress conditions. **(D)** Effect of membrane depolarization on PMT activity. Endogenous PMT activity (MSMEG_5447) was measured following treatment with increasing concentrations of the protonophore CCCP. The untreated WT strain (mc²155) serves as a positive control. High CCCP concentrations (100 µM) result in a loss of detectable activity, concomitant with reduced protein abundance (**Supplementary Figure 9**), whereas moderate concentrations (10 µM) preserve most of the enzymatic activity (∼90%), indicating that PMT function is resilient to partial dissipation of the proton-motive force. Bars represent the mean values ± s.d. of three independent biological replicates.

Similarly, deletion of mprB, htrA, and msmeg_5083 resulted in significant reductions in MtPMT activity (∼65–72% of wild-type), whereas deletion of lppZ had a moderate and non-significant effect (∼80% of wild-type) (**Fig. 6A**). These differential effects likely reflect the distinct roles of these proteins in envelope-associated processes. HtrA, a periplasmic protease, may regulate substrate folding and availability, thereby indirectly modulating O-mannosylation efficiency. MprB, a sensor kinase involved in envelope stress responses, could influence the expression or activity of envelope-associated factors required for optimal MtPMT function. Notably, MSMEG_5083 contains a DivIVA-like domain [59], suggesting a role in membrane organization analogous to the polar scaffold protein Wag31 [60] [61]. Given the established function of DivIVA-family proteins as structural hubs organizing membrane-associated complexes [62], MSMEG_5083 may contribute to the spatial architecture of the MtPMT-associated platform, and its impact on activity likely reflects disruption of this organization rather than direct enzymatic effects.

Importantly, the reduction in MtPMT activity observed upon deletion of these partners is unlikely to result solely from loss of direct physical interactions. Instead, it likely reflects a combination of mechanisms, including (i) altered membrane organization affecting MtPMT localization or stability, (ii) impaired protein translocation limiting substrate access to the glycosylation machinery, and (iii) broader perturbations of envelope proteostasis affecting substrate folding and turnover (**Fig. S9 A**).

Together, these targeted perturbations support the functional relevance of the AP-MS-defined interaction network and suggest that MtPMT operates within a spatially organized membrane platform integrating multiple envelope-associated processes.

Conversely, to further substantiate the physiological relevance of protein O-mannosylation at the cellular level, we assessed the impact of PMT deficiency on bacterial fitness under conditions challenging envelope permeability, structural integrity, stress adaptation, and energetic homeostasis. Given the highly impermeable nature of the mycobacterial envelope [42] [63] [64] [65], we reasoned that defects in its organization should preferentially alter susceptibility to compounds whose entry is limited by barrier properties.

We first evaluated susceptibility to the bulky glycopeptide vancomycin (MW 1449.2 g/mol), which inhibits peptidoglycan biosynthesis but poorly penetrates the intact mycobacterial envelope [63]. While wild-type *M. smegmatis* exhibited only moderate sensitivity, the Δpmt mutant (Δmsmeg_5447) displayed a pronounced increase in susceptibility, particularly at higher concentrations (**Fig. 6B**). This phenotype was fully complemented by reintroduction of MtPMT, demonstrating that the effect is directly attributable to loss of protein O-mannosylation.

We next examined rifampicin (MW 822.9 g/mol), a hydrophobic RNA polymerase inhibitor whose intracellular activity is also limited by envelope permeability. Consistently, the Δpmt strain showed markedly increased sensitivity, further supporting a defect in the envelope barrier restricting entry of bulky compounds.

In contrast, no significant difference in susceptibility was observed with ethambutol (MW 204.31 g/mol), a small-molecule inhibitor of arabinogalactan biosynthesis. This differential phenotype indicates that PMT deficiency does not cause a global collapse of envelope integrity, but instead results in a selective alteration of permeability properties.

We then assessed the role of protein O-mannosylation in stress adaptation. Growth at 42 °C revealed no difference in overall biomass accumulation between wild-type and Δpmt strains. However, the mutant displayed a pronounced and reversible temperature-dependent aggregation phenotype, forming dense clumps resistant to mechanical dispersion, in contrast to homogeneous cultures of the wild-type strain (**Fig. 6C**). This suggests that O-mannosylation modulates the physicochemical properties of the cell surface, thereby influencing cell–cell interactions under stress conditions.

Because the organization of specialized membrane domains in mycobacteria depends on the proton motive force [66], we investigated whether the activity of MtPMT within this membrane platform in influenced by perturbation of membrane energetics. Our interactomic analyses revealed that MtPMT is closely associated with multiple membrane processes (**Fig. 3A, 4B and Fig. S4**) known to depend on the proton-motive force (PMF), including protein translocation [67] [68] [69], respiratory metabolism [69], and cell division [70]. In addition, loss of PMT alters envelope integrity and cell surface properties (**Fig. 6B and C**), while the phage shock protein PspA—a robust MtPMT interactor (**Fig. 2A, 3A, 4B and Table S2**) and O-mannosylated substrate [20] [71]—is a key component of the mycobacterial response to membrane depolarization [72] [73]. Together, these observations prompted us to investigate the functional link between O-mannosylation and membrane homeostasis. Exposure to the membrane-depolarizing protonophore carbonyl cyanide *m*-chlorophenylhydrazone (CCCP), which induces dissipation of the PMF (**Fig. S9 B**), revealed that high concentrations (100µM) induce a global collapse of protein homeostasis (**Fig. S9 C-D**) and loss of detectable O-mannosylation activity (**Fig. 6D**). Whereas moderate depolarization (**Fig. S9 B**) had only a limited effect, with PMT retaining ∼90% of its activity (**Fig. 6D**). The resilience of O-mannosylation to moderate energetic perturbations distinguishes it from many PMF-dependent processes and suggests that POM may represent a previously unrecognized component of the bacterial membrane stress response.

Altogether, these results demonstrate that protein O-mannosylation is embedded within a membrane-associated functional network that integrates envelope permeability, stress adaptation, and energetic homeostasis. This functional organization is consistent with the spatial clustering of MtPMT interactors revealed by AP-MS, supporting the existence of a physiologically relevant membrane platform coordinating multiple envelope-associated pathways.

## Discussion

Understanding how integral membrane enzymes are spatially organized within the bacterial inner membrane remains a fundamental challenge. Here, we introduce a stratified interactomics framework combining multi-stringency AP–MS with a reproducibility-based cumulative persistence score, providing an heuristic scalable strategy to resolve the hierarchical architecture of membrane protein interaction networks. Applied to the *M. tuberculosis* O-mannosyltransferase MtPMT expressed in the *M. smegmatis* surrogate non-pathogenic strain, this approach reveals that bacterial protein O-mannosylation (POM) is not an isolated enzymatic activity but operates at the center of a structured, functionally stratified, and physiologically adaptive membrane platform.

A central and expected finding is the tight spatial integration of MtPMT with the Sec-dependent protein translocation machinery. The consistent co-purification of SecYEG-associated factors, SecDF, YajC, YidC, and FtsH under stringent conditions, corroborated by *in vivo* FRET and AlphaFold3 structural modeling, positions MtPMT within a translocon-associated assembly. This organization provides a mechanistic basis for co- or early post-translocational O-mannosylation, and extends to mycobacteria principles of glycosylation–translocation coupling previously described only in eukaryotes [24] [25]. Critically, it suggests that protein glycosylation in bacteria may be far more spatially constrained than previously assumed — a paradigm shift with implications for our understanding of secreted mannoproteome biogenesis across prokaryotes [54] [58].

Beyond the translocon, the MtPMT-associated platform integrates an unexpected but remarkably coherent set of envelope proteostasis components — membrane proteases, chaperones, and stress-response regulators — whose genetic disruption functionally affects O-mannosylation activity. This interdependence indicates that glycosylation efficiency is not solely determined by enzymatic parameters but depends on the structural and regulatory integrity of the surrounding membrane environment. In this context, membrane-associated chaperones and proteostasis factors may serve as functional surrogates for the MIR domain of eukaryotic PMTs [13] [14], compensating for the absence of a dedicated substrate-recognition module in bacterial enzymes — a mechanistic divergence that warrants direct experimental investigation.

The dynamic remodeling of the MtPMT-associated platform across growth stages adds a temporal dimension to this spatial organization. The transition from a translocation- and synthesis-coupled environment during early stationary phase to a maturation- and stress-oriented assembly at late stationary phase demonstrates that this platform is a physiologically responsive system. Late stationary phase conditions broadly recapitulate features of the nutrient-limited, low-oxygen environment encountered during infection and bacterial persistence within the host. The observed shift toward stress adaptation factors therefore raises the compelling possibility that the MtPMT platform undergoes analogous remodeling during infection, contributing to the known role of POM in *M. tuberculosis* virulence and host immune evasion [30] [31] [32] [33]. Directly characterizing the MtPMT interactome in the native *Mtb* host — particularly under hypoxic or nutrient-limited conditions — represents an essential next step.

The functional diversity and sheer number of MtPMT core interactors argue against a stable stoichiometric complex and instead support a fluid, microdomain-associated platform [5] where repeated transient engagements — facilitated by lateral diffusion within the mycobacterial membrane — enable efficient substrate processing. A striking observation emerging from the comparison with previously reported membrane domain proteomes is that the degree of overlap between the MtPMT interactome identified here and the inner membrane domains (IMDs, formerly called PMf) described in *M. tuberculosis* and *M. smegmatis* by Puffal *et al.* [7] and Hayashi *et al.* [5], respectively, is highly dependent on the bacterial growth stage. Whereas the MtPMT interactome reconstructed at early stationary phase (72 h) shares significant similarity with the membrane proteomes defining the IMD, this overlap becomes markedly reduced within the highly persistent core interactome identified at late stationary phase (96 h) (**Table S3**). Indeed, the 72 h interactome shares 63 and 41 proteins with the Hayashi (437 proteins) and Puffal (304 proteins) datasets respectively, including 38 proteins common to all three datasets. Notably, the extensive overlap observed at 72 h is further reinforced by the presence of several characteristic components of the IMD [5] [7], including Ppm1, PimB, and GlfT2 (**Table S3**), which are central enzymes of cell envelope biosynthesis.

In contrast, these markers are absent from the highly persistent 96 h core interactome, consistent with a substantial remodeling of the membrane environment surrounding MtPMT during bacterial growth. Among the 108 proteins defining the 96 h MtPMT core interactome, only 13 and 9 are shared with the Hayashi and Puffal datasets, respectively, whereas the two IMD proteomes themselves exhibit extensive conservation, sharing 125 proteins despite originating from different mycobacterial species (**Table S3**). Remarkably, only two proteins are common to all three datasets at this stage. Despite this limited overlap, the shared proteins are not randomly distributed but are enriched in structural and functional components involved in respiration, envelope biogenesis and cell division/elongation process (**Table S3**). Among them, the membrane stress protein PspA and the Band-7/flotillin-like protein MSMEG_3155 are particularly noteworthy, as both have been implicated in the organization or stabilization of membrane microdomains [5] [7] [72] [73] [74] [75]. Their recovery within both the MtPMT interactome and IMD-associated proteomes (**Table S3**), supports the idea that MtPMT shares selected organizational principles with previously described membrane domains while defining a distinct functional module rather than representing a generic constituent of the IMD.

These observations highlight the importance of integrating spatial proximity and biochemical robustness as orthogonal parameters to resolve membrane protein interaction architectures. Rather than defining a single interaction map, MtPMT stratified interactomics reveals different layers of dynamic and condition-dependent molecular associations outlining a multi-processes conserved membrane-embedded core. Interestingly, although relatively enriched in components involved cell wall elongation/biogenesis, the composition of the MtPMT core interactome appears to diverge from that of the inner membrane-associated domain previously reported to correlate the polar cell envelope elongation in *M. smegmatis* or *Mtb* [5] [7].

Our findings therefore broaden the current view of membrane compartmentalization in mycobacteria. To date, IMDs have primarily been described as polar and septal membrane subcompartments supporting cell elongation and division [5] [7]. In contrast, our previous super-resolution microscopy analyses demonstrated that MtPMT is distributed throughout the cell periphery rather than being restricted to the poles or septum [18]. Together, these observations raise the possibility that mycobacterial inner membranes contain multiple specialized membrane nanodomains with distinct but coordinated biological functions. The persistent association of MtPMT with the Sec translocon, YidC, FtsY, YajC, FtsH, and other components of the membrane proteostasis network defines a coherent functional platform dedicated to protein secretion, O-mannosylation, membrane protein biogenesis, and quality control. Whether this MtPMT-centered nanodomain operates independently of, or transiently communicates with, the polar IMDs coordinating cell elongation remains an intriguing question. Given the essential contribution of protein O-mannosylation to envelope biogenesis, dynamic coupling between these membrane compartments could provide a mechanism for coordinating protein maturation with bacterial growth and envelope remodeling throughout the bacterial life cycle.

From a therapeutic standpoint, these findings reframe MtPMT as a target embedded within a broader functional network rather than an isolated enzyme. Disrupting not only MtPMT activity but also the spatial integrity of its associated platform — for instance by targeting the MtPMT–FtsH or MtPMT–YajC interfaces — may offer novel avenues for antimycobacterial intervention. The hypersensitivity of *pmt*-deficient strains to bulky antibiotics further suggests that compromising O-mannosylation could synergize with existing envelope-targeting drugs. Similarly, a substantial fraction of MtPMT-associated proteins remains uncharacterized: rather than a limitation, this represents an unexplored reservoir of potentially novel envelope-associated factors whose characterization may uncover new vulnerabilities in the mycobacterial cell surface.

More broadly, the stratified interactomics framework introduced here is not specific to glycosyltransferases or mycobacteria. The challenge of resolving functionally meaningful interactions from membrane-associated background is common to the study of virtually all integral membrane proteins in bacteria. The CPS metric and multi-stringency design provide an example of heuristic strategy for mapping the molecular environment of membrane enzymes across diverse bacterial systems, with potential applications to the study of other medically relevant pathogens.

Collectively, our findings establish that protein O-mannosylation in mycobacteria is intrinsically embedded within a dynamic, hierarchically organized membrane platform that couples glycosylation to protein translocation, envelope proteostasis, and stress adaptation (**Fig. 7**). This model redefines the functional scope of bacterial glycosyltransferases and motivates a new generation of studies aimed at deciphering how spatial organization of the bacterial inner membrane shapes protein maturation, envelope homeostasis, and ultimately, pathogen fitness.

**Figure 7:**
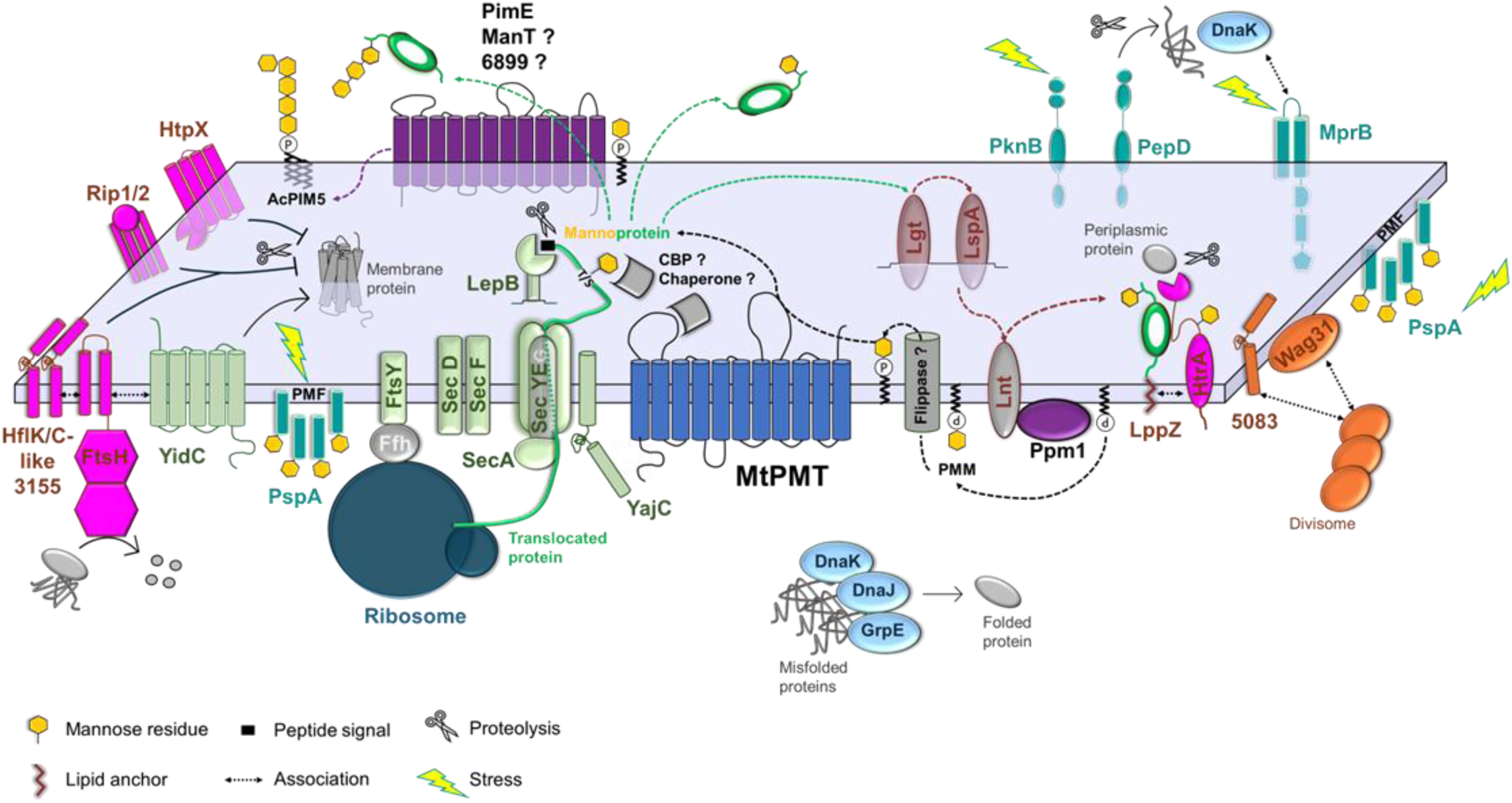
Model of the MtPMT-associated membrane platform integrating protein secretion, glycosylation, proteostasis, and envelope stress responses. Schematic representation of the membrane microdomain in which MtPMT is embedded, illustrating its integration within a dynamic and spatially organized network of envelope-associated processes. MtPMT is positioned in close association with the Sec translocon (green), including SecYEG, SecA, SecDF, YajC, and FtsY, as well as the membrane insertase YidC, supporting co-translocational or early post-translocational *O*-mannosylation. At 72 h, SecA—the ATPase driving post-translational translocation— and ribosomal subunits are localized at the membrane in proximity to the translocon and therefore identified as MtPMT interactors. Ppm1, responsible for the synthesis of the lipid-linked mannose donor (PPM), is also identified as an MtPMT interactor at this stage. In *M. smegmatis*, Ppm1 has been shown to interact with the lipoprotein acylation enzyme Lnt *[2] [3]*, suggesting a functional coupling between *O*-mannosylation and protein acylation processes that may stabilize Ppm1 and enhance *O*-mannosylation efficiency, although Lnt itself is not detected in the MtPMT interactome. PPM is synthesized in the cytoplasm and must be translocated across the membrane to reach the periplasmic side where MtPMT operates. This process likely requires a flippase, although the identity of the PPM flippase remains unknown. The MtPMT interactome may therefore provide a valuable resource to identify candidate proteins involved in this process. Following translocation, substrate proteins are *O*-mannosylated by MtPMT on serine or threonine residues. Signal peptide cleavage is mediated by LepB, which is specifically detected as an interactor at late stationary phase (96 h), consistent with its role in post-translocational maturation. Mannosylated proteins can either reach their final localization or undergo further processing: a substantial fraction is acylated by the lipoprotein maturation system (Lgt, LspA, Lnt; brown/pink), conferring a lipid anchor and leading to the formation of lipomannoproteins associated with the cell envelope. Lgt and LspA are detected as interactors at 96 h. Alternatively, *O*-mannosyl chains can be elongated by others mannosyltransferases (ManT), such as PimE *[4]*, which uses PPM as a donor substrate. PimE is also involved in the biosynthesis of AcPIM5, a major component of the mycobacterial envelope, thereby linking *O*-mannosylation to cell envelope biogenesis (purple). Functional redundancy may exist within this enzyme family, as suggested by the identification of MSMEG_6899, a PimE-like glycosyltransferase, as an MtPMT interactor. In addition, carbohydrate-binding proteins (CBP) and chaperones may facilitate substrate handling by interacting with both MtPMT and its substrates, promoting substrate delivery from the Sec system, stabilizing transient enzyme–substrate interactions, or assisting in substrate release and subsequent targeting to downstream maturation pathways. In parallel, MtPMT is functionally connected to multiple proteostasis systems. Membrane-associated proteases (HtpX, HtrA, Rip1/2; pink), cytoplasmic chaperone machinery DnaK/DnaJ/GrpE (light blue), identified predominantly at 96 h, contribute to protein quality control. FtsH, detected at both 72 h and 96 h, plays a central role in membrane and cytoplasmic protein turnover and is functionally associated with YidC and the Sec machinery *[5]*. In *E. coli*, FtsH activity is regulated by stomatin/prohibitin/flotillin-like proteins (HflK/HflC) *[6] [7] [8]*, and the mycobacterial homolog MSMEG_3155 *[S]*, identified as an MtPMT interactor, may fulfill a similar role. Proteins of this family are known to associate with membrane microdomains *[10] [11] [12] [13]*, suggesting a role in organizing localized membrane environments. HtrA, a mannoprotein, interacts with the lipomannoprotein LppZ *[14]*, which stabilizes its protease activity, contributing to periplasmic protein quality control. Stress sensing and adaptive response systems are also integrated into the MtPMT platform. The two-component system MprB, the chaperone/protease PepD, the kinase PknB, and the phage shock protein PspA (dark bluish green) are identified as interactors, with PspA detected at both 72 h and 96 h. PspA can oligomerize in response to membrane stress and contribute to the maintenance of proton-motive force (PMF) *[15] [16] [17]*. Maintenance of the PMF is essential for multiple processes within this platform, including Sec-dependent protein translocation—particularly via SecDF—YidC *[18] [19] [20]*-mediated membrane protein insertion, and cell division *[21]*. Persistent association of PspA with MtPMT suggests a role beyond a transient substrate, potentially contributing to stabilization of the membrane platform under stress conditions. DnaK may also interact with the periplasmic domain of MprB, contributing to stress sensing and activation of envelope stress responses *[22]*. In addition to MSMEG_3155 and PspA, other scaffolding proteins may contribute to the structural organization of the MtPMT-associated platform, including YajC and DivIVA-like proteins such as Wag31 and MSMEG_5083, which are implicated in membrane organization, divisome assembly, and cell elongation *[23] [24] [25] [26]*. Proteins identified as MtPMT interactors in AP–MS analyses are color-coded according to their functional categories, whereas proteins shown in grey represent functionally associated partners not directly detected or characterized in the interactome. The membrane is represented as a grey bilayer separating the cytoplasm (bottom) from the periplasm (top), with protein topology preserved.

## Materials and Methods

### Reagents and Tools table

**Table 2 :** Strains used in this work.

| Reagent/Resource | Reference or Source | Identifier or Catalog Number |
| --- | --- | --- |
| **Experimental Models** |  |  |
| Strains used in this study | See Table 2 | See Table 2 |
| **Recombinant DNA** |  |  |
| Plasmids used in this study | See Table 3 | See Table 3 |
| **Antibodies** |  |  |
| Anti-m-Cherry | Invitrogen ThermoFisher | MA5-32977 |
| Anti-His | Invitrogen ThermoFisher | 46-0707 |
| Serum Whole Cell Lysate minus LAM Mtb H37Rv | BEI Resources | E293 |
| Goat anti-mouse IgG (H + L), HRP conjugate | Sigma | A4416 |
| Goat anti-rabbit, HRP conjugate | Biorad | 172-1019 |
| **Oligonucleotides and other sequence-based reagents** |  |  |
| Primers used in this study | See Table 4 | See Table 4 |
| **Chemicals, Enzymes and other reagents** |  |  |
| LB broth (Lennox) | Merck | L3022 |
| LB broth with agar (Lennox) | Merck | L2897 |
| Difco Middlebrook 7H11 broth | Becton Dickinson (BD) | 212203 |
| Kanamycin | Sigma-Aldrich | 60615 |
| Hygromycin | Invivogen | Ant-hg1 |
| Streptomycin | Sigma-Aldrich | S9137 |
| Apramycin | Sigma-Aldrich | A2024 |
| Zeocin | Invivogen | ant-zn-1 |
| Vancomycin | Sigma-Aldrich | TA9H97BAEED4 |
| Rifampicin | Sigma-Aldrich | R3501 |
| Ethambutol | Sigma-Aldrich | PHR1930 |
| CCCP (carbonyl cyanide *m*-chlorophenylhydrazone) | Sigma-Aldrich | C2759 |
| DiOC<sub>2</sub>(3) (3,3'-diethyloxacarbocyanine iodide) | Sigma-Aldrich | 320684 |
| Phusion polymerase | Thermo Fisher Scientifics | F530S |
| T4 DNA ligase | Thermo Fisher Scientifics | EL0011 |
| In-Fusion® Snap Assembly Master Mix | Takara | 638948 |
| Low melting-point agarose | Thermo Fisher Scientifics | 16520050 |
| GeneRuler 1 kb DNA ladder | Thermo Scientific | SM0311 |
| NEBuilder HiFi DNA Assembly Cloning Kit | New England Biolabs | E2621S |
| Monarch Plasmid Miniprep Kit | New England Biolabs | T1110L |
| E. coli Stellar chemically competent cells | Takara Bio | 636763 |
| Benzonase | Sigma-Aldrich | E1014-5KU |
| EDTA-free protease inhibitor cocktail | Roche | 04693116001 |
| Digitonin | Sigma-Aldrich | D141 |
| ChromoTek RFP-Trap® Agarose | Proteintech | 17313443 |
| ChromoTek DYKDDDDK Fab-Trap® Agarose (Flag-Trap) | Proteintech | 17872407 |
| 0.2 µm PVDF membranes | Bio-Rad | 1704272 |
| Spectra™ Multicolor Broad Range Protein Ladder | Thermo Fisher Scientifics | 26634 |
| Amersham™ ECL Select™ Western Blotting Detection Reagent | Cytiva | RPN2235 |
| S-Trap | ProtiFi | K002-MINIX-0010KT) |
| Sequencing-grade trypsin | Promega | V5111 |
| UltiMate 3000 system (NCS-3500RS Nano/Cap System) | Thermo Fisher Scientific | 5041.0020 |
| Black clear-bottom 96-well plates | Greiner Bio-One | 655097 |
| 96-well microplates coated with anti-His antibody | Genscript | L00440C |
| Concanavalin A-HRP | Sigma-Aldrich | L6397 |
| TMB solution | eBioscience | 00-4201-56 |
| **Software** |  |  |
| AlphaFold3 | [53] |  |
| Cytoscape 3.10.4 | [76] |  |
| DIA-NN 2.0 | [77] |  |
| Image Lab 3.0 | Bio-Rad |  |
| Mascot 2.7.0 | [78] |  |
| Proline 2.3.3 | [79] |  |
| PyMOL 3.1 | The PyMOL Molecular Graphics System, Version 3.1 Schrödinger, LLC. |  |
| Python 3.14.4 | [80] |  |
| SnapGene | www.snapgene.com |  |
| STRING 12.0 | [81] |  |
| Xcalibur | Thermo Fisher Scientific |  |
| **Other** |  |  |
| CLARIOstar Plus microplate reader | BMG LABTECH |  |
| ChemiDoc Imaging System | Bio-Rad |  |
| Odyssey XF Imaging System | LI-COR Biosciences |  |
| Q Exactive™ Plus Hybrid Quadrupole-Orbitrap™ Mass Spectrometer | Thermo Fisher Scientific |  |
| Orbitrap Exploris™ 480 Mass spectrometer | Thermo Fisher Scientific |  |

| Strains | Genetic backgrounds | Antibiotic resistance | References or sources |
| --- | --- | --- | --- |
| Δpmt | mc<sup>2</sup>155 with its interrupted pmt gene (Δmsmeg_5447) | None | [30] |
| Δpmt::MtPMT | Δpmt complemented by the *Mtb* MtPMT (Rv1002c) protein | Kanamycin | [30] |
| Δpmt::mCherry-MtPMT | Δpmt complemented by the *Mtb* MtPMT (Rv1002c) protein fused at its N-terminus by mCherry (pWM158-N-mCherry plasmid) | Kanamycin | [18] |
| mc<sup>2</sup>155::mCherry-MtPMT | mc<sup>2</sup>155 expressing the *Mtb* MtPMT (Rv1002c) protein fused at its N-terminus by mCherry (pWM158-N-mCherry plasmid) | Kanamycin | This work |
| mc<sup>2</sup>155::Rv1488-mCherry | mc<sup>2</sup>155 expressing the *Mtb* Rv1488 protein fused at its C-terminus by mCherry | Streptomycin | Dr. C. Gutierrez, IPBS |
| mc<sup>2</sup>155::FasC | mc<sup>2</sup>155 expressing the MSMEG_5196 (FasC) protein, tagged with 6xHis | Hygromycin | This work |
| Δpmt::FasC | Δpmt expressing the MSMEG_5196 (FasC) protein, tagged with 6xHis | Hygromycin | This work |
| Δmsmeg_5083 | mc<sup>2</sup>155 with its interrupted msmeg_5083 gene | Zeocin | [51] |
| ΔyajC | mc<sup>2</sup>155 with its interrupted yajC gene (Δmsmeg_2960) | Zeocin | [51] |
| ΔftsH | mc<sup>2</sup>155 with its interrupted ftsH gene (Δmsmeg_6105) | Zeocin | [51] |
| ΔmprB | mc<sup>2</sup>155 with its interrupted mprB gene (Δmsmeg_5487) | Zeocin | [51] |
| ΔhtrA | mc<sup>2</sup>155 with its interrupted htrA gene (Δmsmeg_5070) | Zeocin | [51] |
| ΔlppZ | mc<sup>2</sup>155 with its interrupted lppZ gene (Δmsmeg_2369) | Zeocin | [51] |
| Δmsmeg_5083::FasC | Δmsmeg_5083 expressing the MSMEG_5196 (FasC) protein, tagged with 6xHis | Zeocin, Hygromycin | This work |
| ΔyajC::FasC | Δmsmeg_2960 expressing the MSMEG_5196 (FasC) protein, tagged with 6xHis | Zeocin, Hygromycin | This work |
| ΔftsH::FasC | Δmsmeg_6105 expressing the MSMEG_5196 (FasC) protein, tagged with 6xHis | Zeocin, Hygromycin | This work |
| ΔmprB::FasC | Δmsmeg_5487 expressing the MSMEG_5196 (FasC) protein, tagged with 6xHis | Zeocin, Hygro-mycin | This work |
| ΔhtrA::FasC | Δmsmeg_5070 expressing the MSMEG_5196 (FasC) protein, tagged with 6xHis | Zeocin, Hygro-mycin | This work |
| ΔlppZ::FasC | Δmsmeg_2369 expressing the MSMEG_5196 (FasC) protein, tagged with 6xHis | Zeocin, Hygro-mycin | This work |
| Δpmt::mCherry-MtPMT + YajC-Dendra2-Flag | Δpmt complemented by the *Mtb* MtPMT (Rv1002c) protein fused at its N-terminus by mCherry (pMV366H-N-mCherry plasmid) and expressing YajC protein fused with Dendra2 and Flag at its C-terminal end | Kanamycin, Apramycin | This work |
| Δpmt::mCherry-MtPMT + FtsH-Dendra2-Flag | Δpmt complemented by the *Mtb* MtPMT (Rv1002c) protein fused at its N-terminus by mCherry (pMV366H-N-mCherry plasmid) and expressing FtsH protein fused with Dendra2 and Flag at its C-terminal end | Kanamycin, Apramycin | This work |
| Δpmt::mCherry-MtPMT + MprB-Dendra2-Flag | Δpmt complemented by the *Mtb* MtPMT (Rv1002c) protein fused at its N-terminus by mCherry (pMV366H-N-mCherry plasmid) and expressing MprB protein fused with Dendra2 and Flag at its C-terminal end | Kanamycin, Apramycin | This work |
| Δpmt::mCherry-MtPMT + SodC-Dendra2-Flag | Δpmt complemented by the *Mtb* MtPMT (Rv1002c) protein fused at its N-terminus by mCherry (pMV366H-N-mCherry plasmid) and expressing SodC protein fused with Dendra2 and Flag at its C-terminal end | Kanamycin, Apramycin | This work |
| Δpmt::mCherry-MtPMT + Dendra2-Flag | Δpmt complemented by the *Mtb* MtPMT (Rv1002c) protein fused at its N-terminus by mCherry (pMV366H-N-mCherry plasmid) and expressing the control Dendra2 protein tagged with Flag | Kanamycin, Apramycin | This work |
| Δpmt::Dendra2-Flag | Δpmt expressing the control Dendra2 protein tagged with Flag | Apramycin | This work |

**Table 3:** Plasmids used in this work.

| Plasmids | Description | Antibiotic resistance | References or sources |
| --- | --- | --- | --- |
| pWM158 | Shuttle integrative complementation vector (attP(L5), Ori *E. coli*) carrying the rv1002c gene under the control of the pBlaF* promoter | Kanamycin | [30] |
| pMV366H | Shuttle integrative vector<br>(Ms6, Ori *E. coli*) | Hygro-<br>mycin | [82] |
| pWM218 | Replicative vector carrying the msmeg_5196 gene (fasC), tagged with 6xHis, under the control of the pBlaF* promoter | Hygro-<br>mycin | [30] |
| pWM158-N-mCherry | Plasmid pWM158 with the rv1002c gene fused with mCherry at its N-terminus | Kanamycin | [18] |
| pMV366H-N-mCherry | Integrative pMV366H plasmid carrying the rv1002c gene fused with mCherry at its N-terminus under the control of the pBlaF* promoter | Hygro-<br>mycin | This work |
| pMSR:MSMEG_2960:Dendra | Integrative plasmid carrying the msmeg_2960 (yajC) gene fused with Dendra2 and Flag at its C-terminal end | Apramycin | [51] |
| pMSR:MSMEG_6105:Dendra | Integrative plasmid carrying the msmeg_6105 (ftsH) gene fused with Dendra2 and Flag at its C-terminal end | Apramycin | [51] |
| pMSR:MSMEG_5487:Dendra | Integrative plasmid carrying the Msmeg_5487 (mprB) gene fused with Dendra2 and Flag at its C-terminal end | Apramycin | [51] |
| pMSR:MSMEG_0835:Dendra | Integrative plasmid carrying the Msmeg_0835 (sodC) gene fused with Dendra2 and Flag at its C-terminal end | Apramycin | [51] |
| pMSR:Dendra | Plasmid pMSR:MSMEG_5487:Dendra where the msmeg_5487 gene is deleted, only Dendra2 fused to the Flag tag is retained | Apramycin | This work |

**Table 4:**
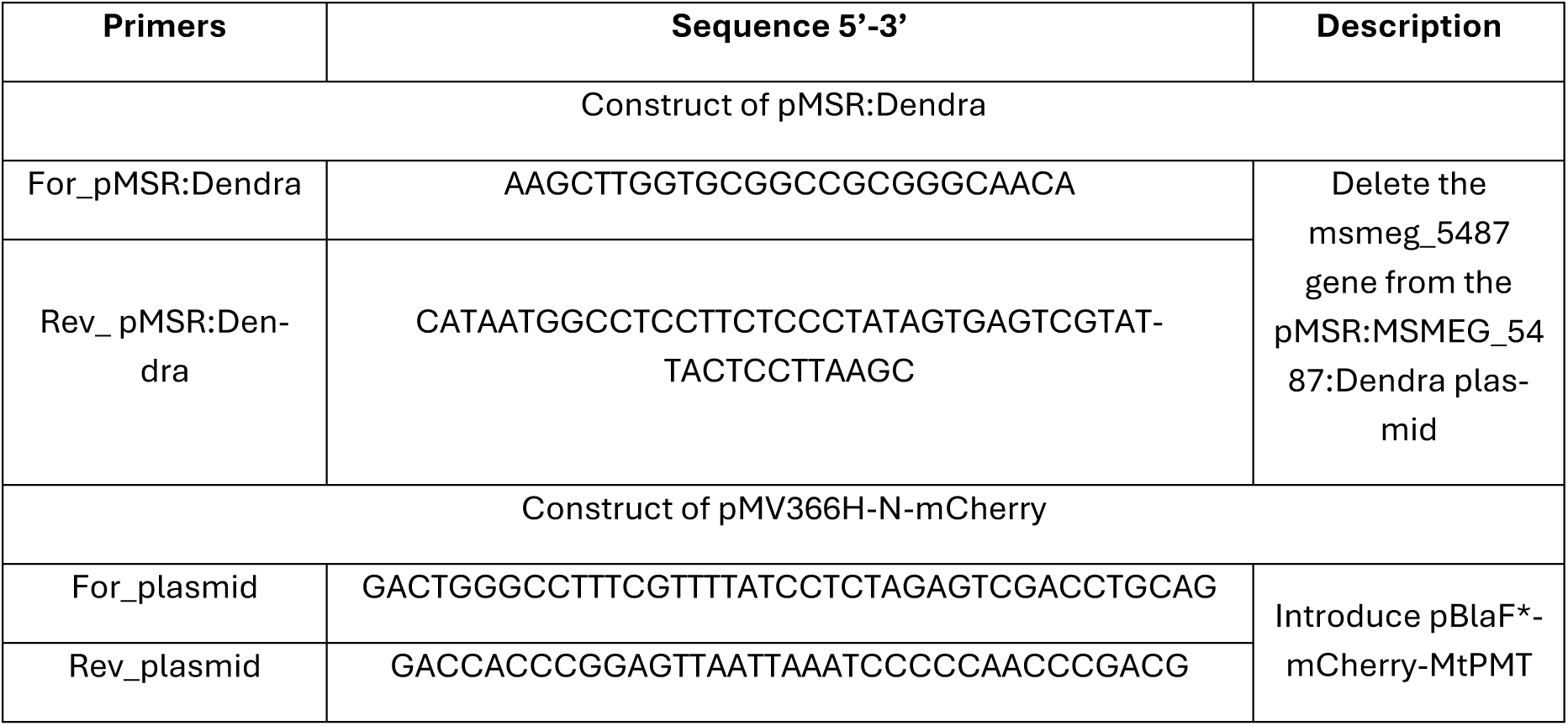
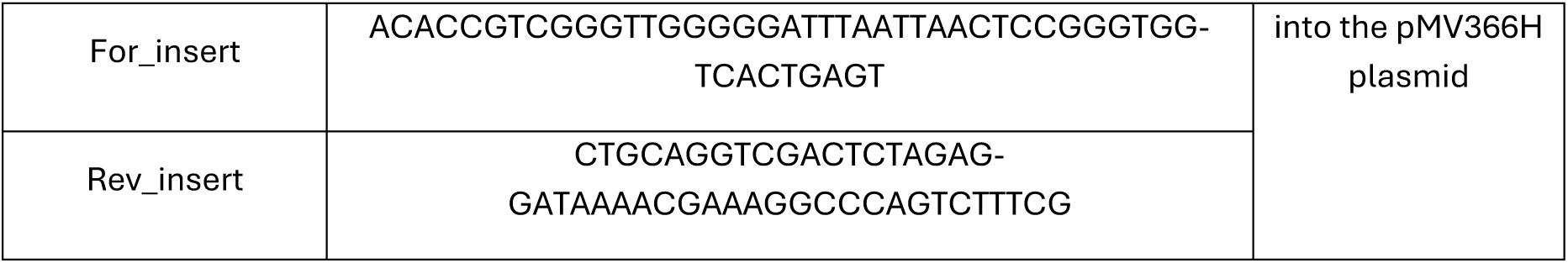
Primers used in this work.

## Methods and Protocols

### Bacterial strains, plasmids and general molecular biology procedures

*Escherichia coli* Stellar chemically competent cells (Takara Bio) were used for plasmid construction, and plasmid amplification. Mycobacterial strains and plasmids used in this study are listed in **Table 2** and **3**. Plasmid maps and sequencing chromatograms were analyzed using SnapGene. Plasmid sequencing was performed by Eurofins. Plasmid DNA was purified using the Monarch Plasmid Miniprep Kit (New England Biolabs).

### DNA constructs

#### Construction of the pMV366H-N-mCherry plasmid

To generate an alternative integrative construct for co-expression experiments, the phage Ms6-derived integrative vector pMV366H was used to express N-terminally tagged mC-MtPMT. The construct was assembled by Gibson assembly (In-Fusion® Snap Assembly Master Mix). DNA fragments were amplified by PCR using Phusion DNA polymerase (Thermo Fisher Scientific), and the fragment containing the *mCherry-MtPMT* fusion under the control of the *pBlaF\** promoter was inserted into pMV366H. Primers used for cloning are listed in **Table 4**. The *mCherry-MtPMT* insert was recovered from previously described plasmids (**Table 3**). The final construct was verified by full plasmid sequencing.

Notably, the MtPMT sequence varies across different databases, with the NCBI Ref.Seq.NP_215518.1featuringashortenedN-terminalendof19 amino acids compared to the UniProt P9WN05 sequence. In this study, we used the shortest version of the enzyme, as the absence of extended N-terminal end does not affect the enzymatic activity [56].

#### Construction of the pMSR:Dendra control plasmid

An integrative plasmid expressing Dendra2-Flag alone, without an additional *M. smegmatis* coding sequence, was generated as a specificity control for reciprocal AP–MS and FRET experiments. Starting from pMSR:MSMEG_5487:Dendra [51], the *msmeg_5487* coding sequence was removed by inverse PCR using Phusion DNA polymerase and outward-facing primers excluding the gene of interest. The final construct was verified by full plasmid sequencing.

### *Mycobacterium Smegmatis* transformation and culture

Plasmids used are listed in Table 3. Electrocompetent *M. smegmatis* cells were prepared from cultures grown in LB broth at 37°C supplemented with 0.05% Tween 80 (LBT). A 10 mL preculture was grown for 7 days to an OD600 of approximately 1.0. Cells were then subcultured into 50 mL fresh LBT and grown to an OD600 of 0.2–0.8. Bacteria were harvested by centrifugation (8 min, 4°C, 500 × g), washed three times with decreasing volumes of ice-cold wash buffer (10% glycerol, 0.05% Tween 80), and resuspended in the final wash buffer. Aliquots of 200 µL were mixed with 1 µL plasmid DNA (100 ng/µL) and electroporated at 2500 V, 25 µF and 200 Ω. Immediately after electroporation, 800 µL LBT were added and cells were allowed to recover for 3 h at 37°C with shaking. Transformants were plated on LB agar containing the appropriate antibiotics and incubated at 37°C for 3–5 days.

Single colonies were grown in 10 mL LBT for 7 days at 37°C to an OD600 of approximately 1.0. When required, kanamycin (25 µg/mL), streptomycin (50 µg/mL), hygromycin (50 µg/mL), apramycin (25 µg/mL), or zeocin (20 µg/mL) were added to the culture media. For interactomic or functional assays, cultures were inoculated at an initial OD600 of 0.05 in larger volumes (50–500 mL). For PMT activity assays, the plasmid expressing FasC-6xHis was introduced into the relevant strains. After 9 days of culture in 10 mL LBT, cells were harvested by centrifugation (10 min, 4°C, 500 × g), and supernatants and pellets were stored separately at −20°C. For reciprocal AP–MS, strains were additionally transformed with pMSR plasmids encoding tagged interactors.

### PMT activity by Enzyme-linked lectin assay (ELLA)

Glycosylation of FasC-6xHis in the culture supernatant was estimated by ELLA as described previously [56]. First, to assess saturable amount of FasC-6xHis for ELLA mannosylation relative quantification, FasC-6xHis concentration from culture supernatants has been quantified by direct-ELISA. For this, wells of a 96 well-plate were filled with 50 μL of culture supernatant, diluted 100× in PBS, and mixed with 50 μL of carbonate coating buffer (Na2CO3 0.53%/ NaHCO 30.42%, pH 9.6) and then incubated overnight at room temperature. After washing 3 times with PBS, the plates were saturated for 1 h at room temperature with 200 μL/well PBS containing 0.05% Tween 20 and 5% milk. After 3 washes with PBS containing 0.05% Tween 20, the plate was incubated for 2 h at room temperature with 50 μL/well of anti-His antibody (anti-His C-term-HRP antibody, Invitrogen), diluted 1000× in PBS containing 0.05% Tween 20 and 0.5% milk. Four washes with PBS were performed before revelation. Subsequently, supernatants with normalized FasC-6xHis concentrations were analyzed for mannosylation by ELLA using 96 well microplates coated with anti-His antibody (Genscript) were used. Plates filled with 100 μL/well of culture supernatants (v/v in PBS) were incubated for 3 h at room temperature and then washed 4 times with PBS containing 2% Tween 20. 100μL of Concanavalin A-HRP solution (Sigma Aldrich) (0.5 μg/mL ConA-HRP, 1mM CaCl2, 1 mM MgCl2, 1mM MnCl2, 0.05% Tween20, in PBS) was added and the plate was incubated at room temperature for 2 h before 4 washes with PBS. Relative values of 100% and 0% were attributed to the amount of ConA binding observed for the WT and *Δpmt M. smegmatis* control strains, respectively. For both assays, revelation was performed with 50 μL/well of TMB solution (eBioscience). 25 μL/well of stop solution (1 mM sulfuric acid) was added after 20 min incubation. Absorbances were measured at 450nm using the BMG LABTECH Clariostar microplate reader.

### PMT activity under CCCP treatment

To assess PMT activity under membrane stress conditions, exponentially growing *M. smegmatis mc²155* cultures expressing FasC-6xHis (OD600 = 0.6) were treated for 24 h at 37°C with 10 µM or 100 µM of CCCP (carbonyl cyanide m-chlorophenyl hydrazone; Sigma). Untreated cultures were used as controls. Cells were harvested by centrifugation (10 min, 4°C, 3000 × g), and supernatants and pellets were stored separately at −20°C. FasC-6xHis abundance and O-mannosylation levels were then measured by direct-ELISA and ELLA, respectively, as described above.

To distinguish direct effects of treatment on PMT activity from the pre-existing O-mannosylation state of FasC-6xHis before treatment, an aliquot of culture supernatant was collected before addition of CCCP. The basal mannosylation signal measured before treatment was subtracted from the relative PMT activity measured after treatment, including in untreated controls.

### Membrane potential measurement

Membrane potential was estimated using the fluorescent probe DiOC₂(3) (3,3′-diethyloxacarbocyanine iodide; Sigma-Aldrich). Bacterial cultures were harvested at an OD600 of 1.0 by centrifugation, and 1 mL aliquots were washed twice with phosphate-buffered saline (PBS). Cell pellets were resuspended in 1 mL of reaction buffer (60 mM Na₂HPO₄, 60 mM NaH₂PO₄, 130 mM NaCl, 5 mM KCl, 0.5 mM MgCl₂, and 10 mM glucose). Aliquots (100 μL) were dispensed in triplicate into black, clear-bottom 96-well microplates (Greiner Bio-One). DiOC₂(3) was added to a final concentration of 30 μM, with or without 5 μM carbonyl cyanide *m*-chlorophenylhydrazone (CCCP; Sigma-Aldrich). Samples were incubated for 35 min at room temperature in the dark. Green and red fluorescence were measured using a CLARIOstar microplate reader (BMG LABTECH) with excitation at 488 nm and emission at 528 nm (green) and 620 nm (red). Membrane potential was estimated from the red-to-green fluorescence ratio (620/528 nm) of DiOC₂(3).

### Antibiotic susceptibility assays on solid medium

Antibiotic susceptibility was assessed by spotting serial dilutions on solid medium. Cultures of *M. smegmatis mc²155*, *Δpmt*, and *Δpmt:MtPMT* were grown in LBT medium supplemented with required antibiotics to OD600 = 1.0. Bacterial suspensions were serially diluted in sterile PBS from 10⁰ to 10^−5^, and 3 µL of each dilution were spotted onto 7H11 agar supplemented with 5% glycerol and vancomycin (0-5 mg/L), or rifampicin (0-10 mg/L), or ethambutol (0-0.3 mg/L). Plates without antibiotics were included as growth controls. Plates were incubated at 37°C for 96 h and growth was visually assessed at 72 h and 96 h.

### Thermal stress assay

Seven-day precultures of *mc²155* and *Δpmt* were inoculated at OD600 = 0.05 into 50 mL fresh LBT medium and incubated at either 37°C or 42°C for 72 h with shaking. Cell density was measured at OD600 and culture morphology was visually examined. Cells were then harvested (5000 rpm, 15 min, 4°C), dried under vacuum, and pellet dry weight was measured. For temperature shift experiments, pellets from cultures initially grown at 37°C were resuspended in 50 mL fresh LBT and incubated at 42°C for 12 h, whereas pellets from cultures initially grown at 42°C were resuspended in fresh LBT and incubated at 37°C for 12 h. Culture morphology was then reassessed.

### Whole-cell fluorescence measurements

For fluorescence intensity measurements, 1 mL bacterial cultures were harvested by centrifugation (10 min, 5000 rpm), washed three times in PBS and resuspended in the same volume. Cells were transferred in triplicate to black clear-bottom 96-well plates (Greiner Bio-One). mCherry fluorescence was measured at 610 nm after excitation at 587 nm, and Dendra2 fluorescence at 507 nm after excitation at 490 nm, using a CLARIOstar microplate reader. Optical density at 600 nm was used for normalization.

### FRET analysis in live bacteria

Interactions between MtPMT and selected candidate partners were further analyzed by Förster resonance energy transfer (FRET) using whole bacterial cells expressing the corresponding fusion proteins. Constructs encoding mC-MtPMT and Dendra2-tagged proteins (YajC-Dendra2-Flag, FtsH-Dendra2-Flag, MprB-Dendra2-Flag, SodC-Dendra2-Flag, or Dendra2-Flag alone) were introduced into *M. smegmatis mc²155* by electroporation. Transformed strains were grown in LBT medium containing the appropriate antibiotics for 120 h. Expression of fusion proteins was verified by whole-cell fluorescence measurements and western blotting.

For FRET measurements, cells were harvested (5000 × g, 15 min, 4°C), washed twice in PBS and resuspended in the same buffer. Black clear-bottom 96-well plates were loaded with 200 µL samples in triplicate. FRET was measured using a CLARIOstar plate reader by exciting the donor fluorophore Dendra2 at 490 nm and measuring emission at 650 nm corresponding to mCherry.

Corrected FRET intensity was calculated as:

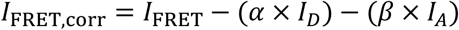

where *I_D_* is donor fluorescence leakage into the FRET channel, *I_A_*is direct excitation of the acceptor in the FRET channel, *_α_*is the donor bleed-through coefficient, and *β* is the acceptor direct excitation coefficient. Control strains expressing Dendra2-Flag alone or mC-MtPMT alone were used to calculate these coefficients. Corrected FRET values were normalized to cell density (OD600).

The SodC-Dendra2-Flag control was used as a baseline for non-interacting proteins.

### Cell lysis and membrane protein extraction

*M. smegmatis*, cultures expressing mCherry-MtPMT, Dendra2-Flag constructs or controls were harvested after 72 h or 96 h of growth (**Table 5**). Unless otherwise stated, all steps were performed on ice or at 4°C.

**Table 5:**
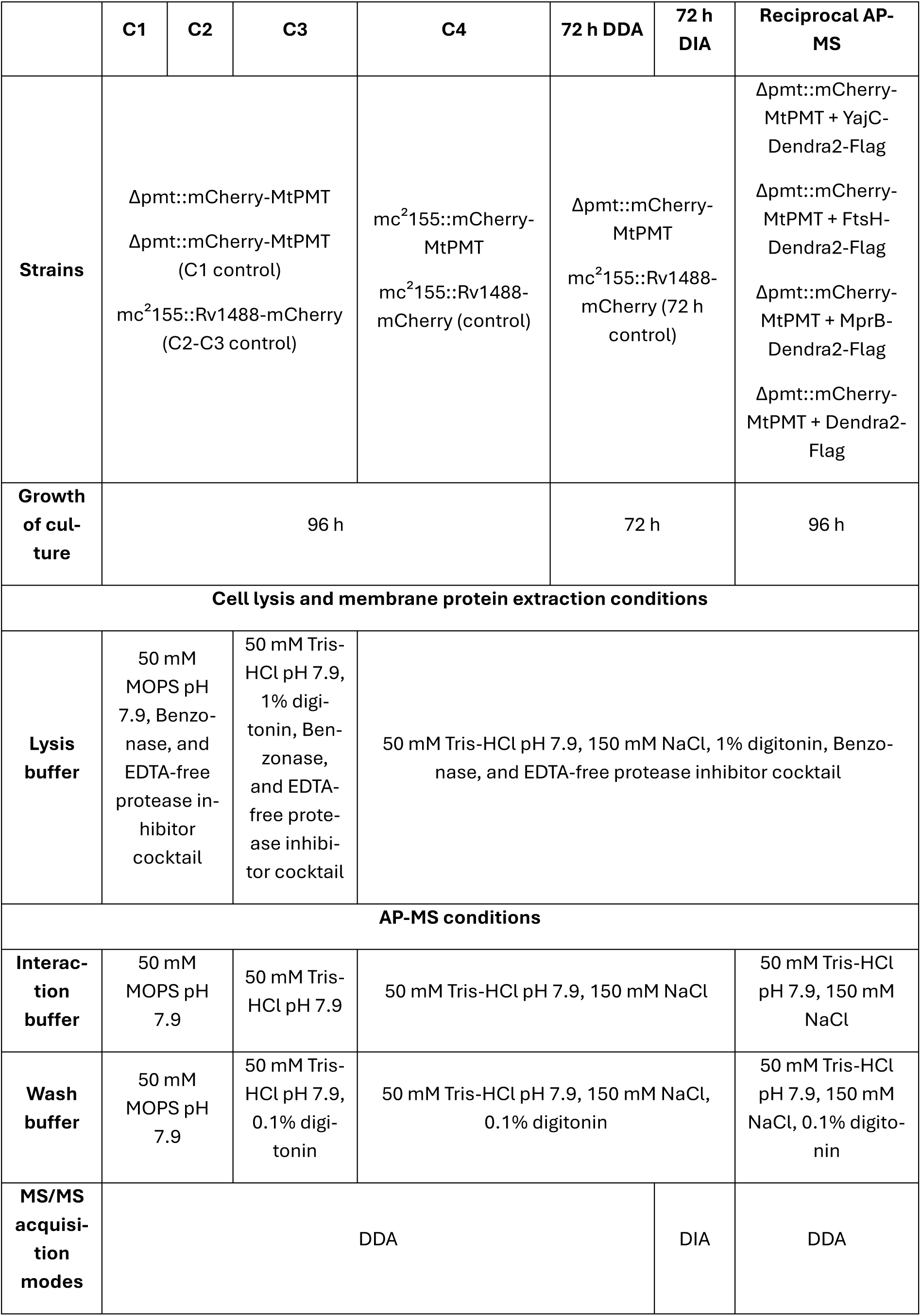
Cell lysis, membrane protein extraction and AP-MS experimental design.

400 mL cultures grown in LB containing 0.05% Tween 80 and the appropriate antibiotics were harvested. Cell pellets were frozen at −20°C to weaken the cell wall, centrifuged (15 min, 5000 rpm, 4°C), and washed twice with cold PBS.

Cells were resuspended in lysis buffer (**Table 5**) using 1 mL buffer per mg wet pellet. Mechanical lysis was performed with zirconium beads using a bead beater (4 cycles of 1 min, separated by 2 min cooling on ice). Lysates were clarified by centrifugation (3000 × g, 10 min, 4°C) to remove unbroken cells and debris, incubated for 4 h at 4°C under gentle agitation to extract membrane proteins, and centrifuged again (13,000 × g, 10 min, 4°C) to remove insoluble material. Protein concentration in the soluble fraction was estimated by absorbance at 280 nm.

Extraction efficiency of mCherry–MtPMT was systematically monitored by western blot using anti-mCherry antibodies.

### SDS–PAGE and western blotting

Whole-cell lysates were separated by SDS–PAGE. Samples (12 µL) were mixed with 3 µL Laemmli buffer, incubated for 15 min at 50°C, and loaded onto gels. After migration, gels were transferred to nitrocellulose membranes for western blot analysis.

For immunodetection, membranes were incubated with the appropriate primary antibody diluted in PBS containing 1% BSA and 0.05% Tween 20 for 3 h, blocked in PBS containing 5% BSA and 0.1% Tween 20 for 2 h, and then incubated with the appropriate secondary antibody diluted in PBS containing 1% BSA and 0.1% Tween 20 for 1 h. Detection was performed using Amersham ECL Prime according to the manufacturer’s instructions.

### Analysis of mC-MtPMT expression after CCCP treatment

To assess the effect of membrane stressors on MtPMT abundance, the *Δpmt* strain complemented with mC-MtPMT was grown from 7-day precultures diluted to OD600 = 0.05 in 50 mL fresh LBT medium containing the appropriate antibiotic selection. Once cultures reached OD600 = 0.6, they were treated for 24 h at 37°C with 10 µM or 100 µM of CCCP. Untreated cultures served as controls.

Cells were lysed (lysis buffer: 50 mM Tris-HCl pH 7.9, 150 mM NaCl, 1% digitonin, Benzonase, and EDTA-free protease inhibitor cocktail) as described above, and total protein concentration was estimated by absorbance at 280 nm. Expression of mC-MtPMT was analyzed by western blot using anti-mCherry antibodies (Invitrogen), and anti-*M. tuberculosis* serum (Serum Whole Cell Lysate minus LAM Mtb H37Rv, BEI Resources) was used as a loading normalization control.

### Affinity purification of mCherry-MtPMT

Affinity purification of mCherry-MtPMT and associated complexes was performed using RFP-Trap Agarose beads (Chromotek) according to the manufacturer’s instructions. Briefly, 500 µL clarified lysate were diluted 1:1 in interaction buffer lacking non-ionic detergent (**Table 5**) and incubated overnight at 4°C with 50 µL pre-washed bead slurry. Under these conditions, capture was performed in the presence of 0.5% digitonin (except for C1-C2). Beads were collected by gentle centrifugation (3000 × g, 5 min, 4°C) and washed three times with wash buffer (**Table 5**). Bound proteins were eluted with 5% SDS at 50°C for 15 min, followed by a second elution in 5% SDS at 95°C for 5 min. Eluates were stored at −20°C until proteomic analysis. Purification efficiency was monitored by western blot.

### Affinity purification of Flag-tagged proteins complexes

*M. smegmatis mc²155* strains co-expressing mC-MtPMT and YajC-Dendra2-Flag, FtsH-Dendra2-Flag, MprB-Dendra2-Flag or Dendra2-Flag alone were grown for 96 h. Expression of fluorescent proteins was verified by whole-cell spectrofluorimetry and/or western blotting prior to lysis.

Lysates were prepared as described above (**Table 5**) and subjected to affinity purification using DYKDDDDK Fab-Trap Agarose beads (Chromotek) according to the manufacturer’s recommendations. Briefly, 500 µL lysate were diluted 1:1 in detergent-free interaction buffer (**Table 5**) and incubated overnight at 4°C with 50 µL pre-washed bead slurry. Beads were washed three times in wash buffer (**Table 5**), and bound proteins were eluted with 5% SDS at 50°C. Eluates were analyzed by western blot using anti-mCherry antibodies to assess co-purification of mC-MtPMT with the candidate Flag-tagged partners.

### Proteomics analyses

#### Peptide preparation for mass spectrometry

Protein samples were processed using the S-Trap protocol (Protifi). Proteins were reduced with 10 mM DTT (10 min, 95°C), alkylated with 20 mM iodoacetamide (30 min, room temperature, protected from light), acidified with phosphoric acid (final concentration 1.2%), and diluted in binding buffer containing 90% methanol in 100 mM Tris-HCl pH 7.1. Protein precipitates were captured on mini S-Trap columns, washed according to the manufacturer’s instructions, and digested on-column with sequencing-grade trypsin (Promega) in 50 mM TEAB pH 8.0 at an enzyme-to-substrate ratio of 1:25 (w/w) for 16 h at 37°C. Peptides were sequentially eluted with 50 mM TEAB, 0.2% formic acid, and 50% acetonitrile/0.2% formic acid. Eluates were pooled, dried in a SpeedVac, and stored at −20°C.

#### NanoLC–MS/MS analysis in DDA mode

Dried peptide samples were resuspended in solvent A (5% acetonitrile, 0.05% trifluoroacetic acid), and 5 µL were injected for nanoLC–MS/MS analysis. Peptides were loaded onto a C18 precolumn (300 µm ID × 5 mm, Dionex) at 20 µL/min, desalted for 5 min, and separated on a PepMap C18 analytical column (75 µm ID × 15 cm, Dionex) using a gradient from 5% to 50% solvent B (80% acetonitrile, 0.2% formic acid) at 300 nL/min.

For data-dependent acquisition (DDA), spectra were acquired on a Q Exactive Plus instrument (Thermo Fisher Scientific). MS1 scans were acquired in the Orbitrap at 70,000 resolution over a 350–1600 m/z range. The 15 most intense ions were selected for HCD fragmentation (normalized collision energy 28%), and MS2 spectra were acquired at 17,500 resolution. Dynamic exclusion was set to 30 s.

#### Nano LC-MS/MS analysis in DIA mode

Tryptic peptides were resuspended in 50 µl of 2% acetonitrile and 0.05% trifluoroacetic acid and analyzed by nano-liquid chromatography (LC) coupled to tandem MS, using an UltiMate 3000 system (NCS-3500RS Nano/Cap System; Thermo Fisher Scientific) coupled to an Orbitrap Exploris 480 mass spectrometer (Thermo Fisher Scientific). One microliter of each sample was loaded on an analytical C18 column (PEPMAP C18 2 μm 75 μm × 500 mm Thermo Fisher Electron) heated at 45 ◦C, equilibrated in 95% solvent A (5% acetonitrile, 0.2% formic acid) and 5% solvent B (80% acetonitrile, 0.2% formic acid). Peptides were eluted using a 95 min gradient: 0–25% buffer B for 58 min, 25–40% buffer B for 20 min, 40–90% buffer B for 2 min, and 90% buffer B for 5 min. The mass spectrometer was operated in data independent acquisition (DIA) mode with the Xcalibur software. Full-scan MS resolution was set at 60,000, and the AGC target was 300% with a maximum injection time of 100 ms. The MS1 mass range was set to 400–1008. For MS2 spectra, AGC target value spectra were set at 1000%. The DIA isolation scheme consisted of 2 × 75 staggered windows (8 m/z width, 4 m/z offset) covering the 400–1000 m/z range. The resolution was set to 15,000, injection time was set in automatic mode, and normalized collision energy was set at 30%. Raw data were processed using DIA-NN (version 2.0) using default parameters and the spectral library generated with the DDA analysis. DIA-NN reports were postprocessed using the Proline software v2.3.3.

#### Proteomic data processing and quantification

Label-free quantification was performed using Proline by extracting and integrating chromatographic peak intensities of proteotypic peptides. Intensities were normalized across samples to correct for experimental and instrumental variability.

Raw MS files were processed with the Mascot software (version 2.7.0) for database search and Proline for label-free quantitative analysis (version 2.3.3). Data were searched against *Mycobacterium smegmatis* (strain ATCC 700084 / mc²155), 6,603 entries of the UniProtKB protein database supplemented with the sequence of mCherry–MtPMT construct. Carbamidomethylation of cysteines was set as a fixed modification, while methionine oxidation, deamidation of asparagine, and hexose modifications on serine/threonine residues were considered as variable modifications. Specificity of trypsin/P digestion was set for cleavage after K or R, and two missed trypsin cleavage sites were allowed. The mass tolerance was set to 10 ppm for the precursor and to 20 mmu in tandem MS mode. Minimum peptide length was set to 7 amino acids, and identification results were further validated in Proline by the target decoy approach using a reverse database at both a PSM and protein false-discovery rate of 1%. For label-free relative quantification of the proteins across biological replicates and conditions, cross-assignment of peptide ions peaks was enabled inside group with a match time window of 1 min, after alignment of the runs with a tolerance of +/- 600s.

Median Ratio Fitting computes a matrix of abundance ratios calculated between any two runs from ion abundances for each protein. For each pair-wise ratio, the median of the ion ratios is then calculated and used to represent the protein ratio between these two runs. A least-squares regression is then performed to approximate the relative abundance of the protein in each run in the dataset. This abundance is finally rescaled to the sum of the ion abundances across runs. A Student t-test (two-tailed t-test, equal variances) was then performed on log2 transformed values to analyze differences in protein abundance in all biologic group comparisons. Significance level was set at p<0.05, and ratios were considered relevant if higher than +/- 1.

The mass spectrometry proteomics data have been deposited to the ProteomeXchange Consortium via the MassIVE partner repository with the dataset identifier MSV000103135.

PCA was performed on normalized abundance values obtained from all quantified proteins identified across AP–MS conditions (C1–C4). Proteins detected in fewer than 70% of samples were excluded prior to analysis. Missing values were imputed using the minimum abundance value observed for each protein, followed by log₂ transformation of abundance values. Principal component analysis was subsequently performed on standardized quantitative profiles using the first two principal components for visualization. Replicate samples were analyzed independently and visualized according to AP–MS purification condition.

Hierarchical clustering analysis was performed to compare protein enrichment profiles across AP–MS purification conditions. Proteins displaying significant enrichment or depletion (|log2 fold change| ≥ 1 and *p* ≤ 0.05) in at least one condition were retained for analysis. Missing values were replaced by zero after merging datasets across purification conditions. Clustering was performed using Euclidean distance and average linkage on log2 fold change values using the Python package Seaborn (v3.14.4) implemented through the clustermap function.

### Interactome stratification and scoring

To assess interaction robustness, proteins were classified based on their detection across four purification conditions (C1-C4) of increasing stringency. Interactors were grouped into four classes: core (detected in all conditions), stable (three conditions), conditional (two conditions), and peripheral (one condition).

To quantitatively capture interaction persistence across conditions, a persistence score was assigned to each protein based on its presence in each dataset. Detection in C1, C2, C3, and C4 was assigned a value of 1, 2, 3, and 4, respectively. The cumulative persistence score (CPS) was then calculated as the sum of these values across all conditions in which the protein was detected.

### Functional enrichment analysis

Functional enrichment analyses were performed using KEGG pathway [48] and Gene Ontology (GO) [49] databases. Enrichment significance was calculated using appropriate statistical tests with FDR correction.

### Network analysis

Protein interaction networks were explored using the STRING database [81], integrating evidence from co-expression, co-occurrence, and experimental datasets. The resulting networks were imported into Cytoscape [76] for visualization and graphical representation.

### Structural modeling

Structural models were generated using AlphaFold3-based predictions [53]. To investigate the spatial organization of MtPMT within its membrane-associated interaction network, MtPMT was modeled in complex with components of the Sec-dependent protein translocation machinery and associated factors.

Specifically, MtPMT was modeled together with the core Sec holotranslocon components SecY, SecE, SecG, SecD, SecF, YajC and YidC [54] [58]. In addition, Sec-associated proteins identified in the MtPMT interactome by AP–MS (96 h) were included in the modeling, namely the signal recognition particle receptor FtsY, the signal peptidase LepB, and the membrane protease FtsH, which is functionally linked to the Sec translocation system [55].

All protein sequences used for structural predictions correspond to *Mycobacterium tuberculosis* homologs and were retrieved from the UniProt accession number (MtPMT: P9WN05; YajC: P9WL75; FtsH: P9WQN3; SecE: P9WGN7; SecG: P9WGN5; SecY: P0A5Z2; SecD: Q50634; SecF: P9WGN9; YidC: P65626; FtsY: P66842; LepB: Q10789).

Predicted structures were analyzed and visualized using PyMOL.

### Statistical analysis

Unless otherwise stated, bar charts and scatter (XY) plots represent independent biological replicates, defined as independent bacterial cultures derived from distinct clones. Error bars correspond to the standard deviation (s.d.) when *n* ≥ 3.

For comparisons of enrichment score distributions across AP–MS conditions for the 108 common interactors (**Fig. 1G**), paired two-sided Wilcoxon signed-rank tests (scipy.stats.wilcoxon) were performed. Because the same set of proteins was analyzed across all conditions, a paired non-parametric framework was used. Resulting *P*-values were adjusted for multiple comparisons using the Bonferroni correction. Violin plots display individual values together with the median and interquartile range.

For FRET analyses (**Fig. 5D**), each bacterial strain was analyzed using three independent biological replicates, each comprising three technical replicates. Technical replicates were averaged within each biological replicate, resulting in independent values per strain. Individual replicate values are displayed in the graphs. Statistical comparisons were performed against the negative control (mC-MtPMT + SodC–Dendra2–Flag) using unpaired two-tailed Welch’s *t*-tests (scipy.stats.ttest_ind, equal_var=False).

For PMT activity measurements (**Fig. 6A**), each strain (“WT”, “Δpmt”, “Δ5083”, “ΔmprB”, “ΔlppZ”, “ΔhtrA”, “ΔftsH”, and “ΔyajC”) was analyzed using four independent biological replicates, each comprising three technical replicates. Technical replicates were averaged within each biological replicate prior to analysis. Individual replicate values are displayed in the graphs. Statistical comparisons were performed between the WT strain and each mutant using unpaired two-tailed Welch’s *t*-tests (scipy.stats.ttest_ind, equal_var=False).

For CCCP treatment experiments (**Fig. 6D**), three independent biological replicates, each comprising three technical replicates, were analyzed. Technical replicates were averaged within each biological replicate, and results are reported descriptively without formal statistical testing.

Unless otherwise stated, no correction for multiple comparisons was applied. Statistical significance was defined as ns, not significant; * *P* < 0.05; ** *P* < 0.01; *** *P* < 0.001; **** *P* < 0.0001.

All statistical analyses were performed using Python (SciPy library, v1.17.1).

## Data availability

The mass spectrometry datasets generated during and analyzed during the current study are available in the MassIVE repository (MSV000103135) and are available from the corresponding author upon reasonable request.

## Acknowledgments

We express our gratitude to Claude Gutierrez (IPBS, Toulouse, France) for providing the mCherry-Rv1488 plasmid used for pull-down control experiments.

## Author contributions

All authors contributed extensively to the work presented in this paper. C.R. conducted all the experiments. G.P. contributed to the preanalytical experimental steps. A.S. performed the MS analyses under O.S. supervision. J.N. gave technical support and conceptual advice. M.R. designed the project and supervised the experiment. CR and MR wrote the manuscript. The authors used an AI-based language model to assist with English editing and clarity. All scientific content, interpretations, and conclusions were developed and validated by the authors.

---

**Supplementary Figure 1:**
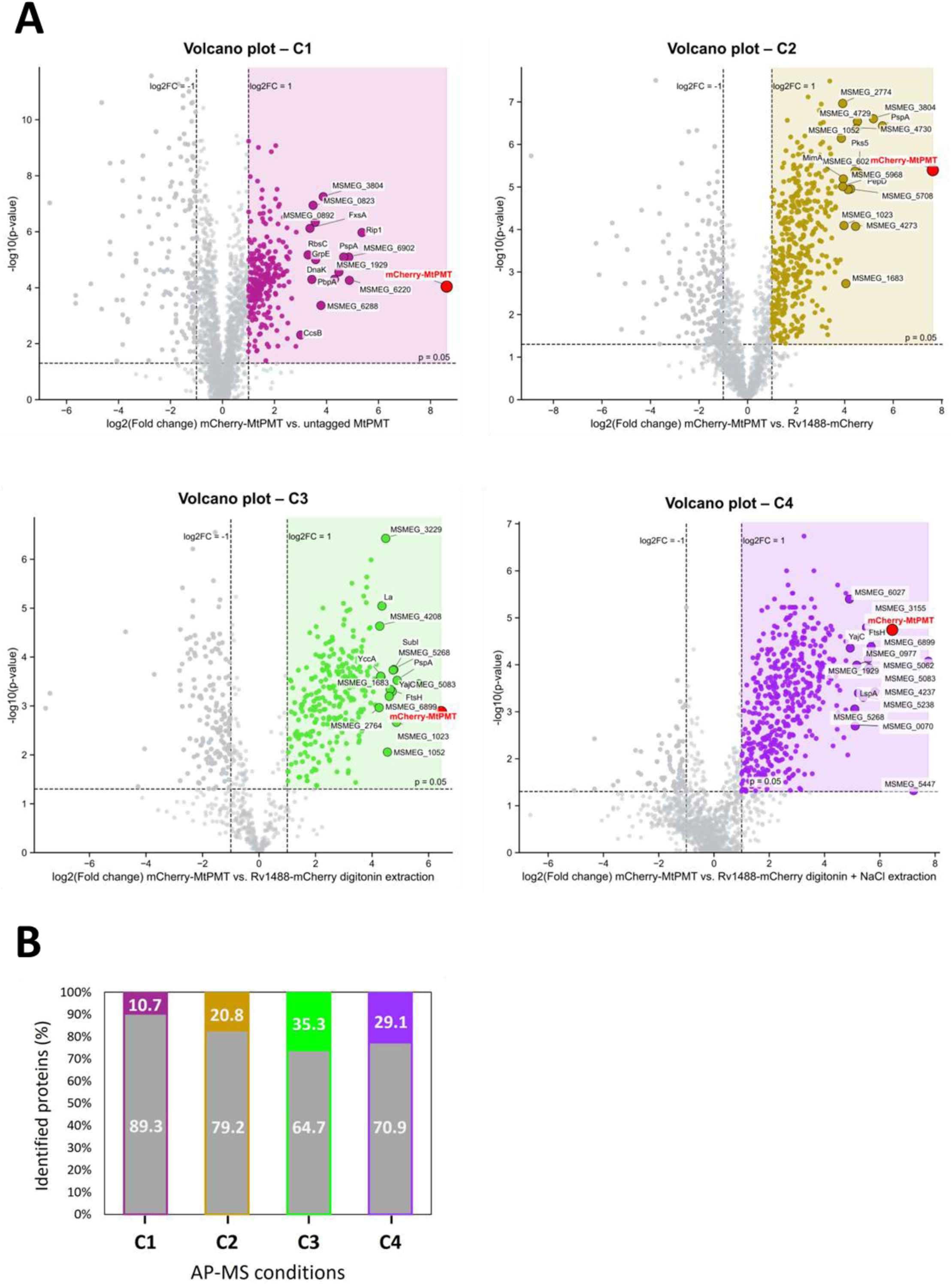
Progressive enrichment and selectivity of MtPMT interactors across AP–MS conditions. **(A)** Volcano plots corresponding to the four AP–MS conditions (C1–C4). For each condition, protein enrichment is represented as log₂(fold change) versus –log₁₀(P-value), comparing mCherry–MtPMT pulldowns to the appropriate control. Proteins passing the selection thresholds (log₂FC ≥ 1 and P < 0.05) are highlighted in color and represent candidate MtPMT interactors, whereas non-specific proteins are shown in grey. The 15 most enriched candidate interactors are annotated in each plot and are listed in **Supplementary Table 1**. The bait protein mCherry–MtPMT is indicated in red. **(B)** Proportion of candidate interactors across AP–MS conditions. Colored fractions correspond to candidate interactors, whereas grey fractions represent non-selective proteins. Progressive refinement of purification conditions leads to an increase in the proportion of selective interactors, with maximal enrichment observed under digitonin extraction (C3) and sustained selectivity under high-stringency conditions (C4).

**Supplementary Table 1:** Comparison of the top 15 enriched candidate interactors across AP– MS conditions. Interactors are color-coded according to the purification condition: magenta for C1 (untagged MtPMT control, detergent-free), yellow for C2 (Rv1488–mCherry control, detergent-free), green for C3 (digitonin extraction), and purple for C4 (digitonin combined with high ionic strength, NaCl). For each dataset, proteins are ranked based on their log₂ fold change relative to the control condition.

| Gene name | Alternative name | Localisation | Description | Log2(FC) | -Log10 (P-value) |
| --- | --- | --- | --- | --- | --- |
| **C1** |  |  |  |  |  |
| **rip1** |  | Plasma membrane | Zinc metalloprotease Rip1, Regulated in-tramembrane proteolysis during stress | 5.37 | 5.97 |
| **MSMEG_6220** |  | Outer membrane | Lipoprotein | 4.88 | 4.26 |
| **MSMEG_6902** |  | Plasma menbrane | Putative membrane protein | 4.86 | 5.09 |
| **MSMEG_2695** | **35 kDa / PspA** | Cytoplasm,<br>Plasma membrane | 35 kDa protein, Phage shock protein (HSP70 family) | 4.66 | 5.09 |
| **MSMEG_1929** |  | Plasma membrane | Conserved alanine valine rich protein | 4.48 | 4.56 |
| **pbpA** |  | Plasma membrane | Penicillin-binding protein A, Peptidoglycan biosynthetic process, Regulation of cell shape | 4.31 | 4.30 |
| **MSMEG_3804** |  |  | Phage shock protein A (IM30), an orthologue of 35 kDa protein | 3.88 | 7.25 |
| **MSMEG_6288** | **LucA** | Plasma membrane | Lipid uptake coordinator A, required for full virulence | 3.79 | 3.37 |
| **grpE** |  | Cytoplasm | Co-chaperone GrpE | 3.58 | 5.00 |
| **MSMEG_0892** |  | Membrane | Uncharacterized protein | 3.57 | 6.34 |
| **MSMEG_0823** |  | Membrane | Septum_form domain-containing protein | 3.49 | 6.94 |
| **dnaK** |  | Plasma membrane periphery, Cytoplasm,<br>Periplasm,<br>Extracellular region | Chaperone protein DnaK (HSP70) | 3.44 | 4.29 |
| **MSMEG_3862** | **FxsA** | Plasma membrane | FxsA cytoplasmic membrane protein | 3.37 | 6.12 |
| **MSMEG_6803** | **RbsC** | Plasma membrane | Ribose transport system permease protein RbsC | 3.29 | 5.17 |
| **ccsB** |  | Plasma membrane | Cytochrome c-type biogenesis protein CcsB | 3.01 | 2.31 |
| **C2** |  |  |  |  |  |
| **MSMEG_2695** | **35 kDa / PspA** | Cytoplasm,<br>Plasma membrane | 35 kDa protein, Phage shock protein (HSP70 family) | 5.56 | 6.44 |
| **MSMEG_3804** |  |  | Phage shock protein A (IM30), an orthologue of 35 kDa protein | 5.19 | 6.60 |
| **pks5** |  | Membrane | Mycocerosic acid synthase-like polyketide synthase | 4.57 | 5.33 |
| **MSMEG_4729** |  |  | Fatty acyl-AMP ligase FadD28 and polyketide synthase | 4.53 | 6.54 |
| MSMEG_4730 |  |  | Fatty acyl-AMP ligase FadD28 and polyketide synthase | 4.48 | 6.37 |
| MSMEG_4273 |  | Plasma membrane | Integral membrane protein | 4.45 | 4.07 |
| MSMEG_6027 |  | Plasma membrane | Major facilitator superfamily protein MFS_1 | 4.43 | 5.37 |
| MSMEG_5486 | PepD | Plasma membrane, Extracellular region | Peptidase S1 and S6, chymotrypsin/Hap, serine protease PepD, Proteolysis, cleaves several proteins, including the 35 kDa antigen PspA, Plays an important role in the stress response network mediated through the two-component regulatory system MprAB and SigE signaling networks, Required for virulence | 4.24 | 4.95 |
| MSMEG_5708 |  | Membrane | DUF2510 domain-containing protein | 4.14 | 4.94 |
| MSMEG_1683 |  | Plasma membrane | Cytosine/purine/uracil/thiamine/allantoin permease family protein | 4.05 | 2.73 |
| MSMEG_1023 |  | Membrane | Integral membrane transporter | 3.97 | 4.09 |
| mimA |  |  | Propane 2-monooxygenase, hydroxylase component large subunit | 3.94 | 5.19 |
| MSMEG_5968 |  | Plasma membrane | Polysaccharide biosynthesis protein | 3.93 | 5.01 |
| MSMEG_2774 |  | Membrane | Diguanylate cyclase (Ggdef) domain protein | 3.92 | 6.96 |
| MSMEG_1052 |  | Plasma membrane | Amino acid carrier protein, sodium/alanine symporter | 3.87 | 6.14 |
| C3 |  |  |  |  |  |
| MSMEG_5083 |  | Membrane | DivIVA domain-containing protein, Cell division protein | 5.66 | 3.45 |
| MSMEG_2695 | 35 kDa / PspA | Cytoplasm, Plasma membrane | 35 kDa protein, Phage shock protein (HSP70 family) | 4.89 | 3.52 |
| MSMEG_1023 |  | Membrane | Integral membrane transporter | 4.86 | 2.67 |
| MSMEG_5268 |  | Plasma membrane | Probable proline and glycine rich transmembrane protein | 4.77 | 3.74 |
| MSMEG_4533 | SubI | Outer membrane-bounded periplasmic space | Sulfate-binding lipoprotein SubI, ABC transporter | 4.75 | 3.74 |
| ftsH |  | Plasma membrane | ATP-dependent zinc metalloprotease FtsH, Cell division protein, Membrane-bound protease, Plays a role in the quality control of integral membrane proteins | 4.71 | 3.30 |
| yajC |  | Plasma membrane | Sec translocon accessory complex subunit YajC | 4.64 | 3.34 |
| MSMEG_6899 |  | Plasma membrane | Predicted to be in the GT-C superfamily of glycosyltransferases, glycosyltransferase 87 family | 4.61 | 3.20 |
| MSMEG_1052 |  | Plasma membrane | Amino acid carrier protein, sodium/alanine symporter | 4.55 | 2.06 |
| MSMEG_3229 |  | Membrane | DUF4190 domain-containing protein | 4.48 | 6.43 |
| MSMEG_1958 | La | Extracellular region | Endopeptidase La | 4.35 | 5.04 |
| MSMEG_1683 |  | Plasma membrane | Cytosine/purine/uracil/thiamine/allantoin permease family protein | 4.31 | 3.60 |
| MSMEG_5275 | YccA | Membrane | Permease of the major facilitator superfamily protein | 4.30 | 3.61 |
| MSMEG_4208 |  | Membrane | Integral membrane protein | 4.27 | 4.63 |
| MSMEG_2764 |  | Membrane | Probable conserved alanine rich transmembrane protein | 4.25 | 2.97 |
| **C4** |  |  |  |  |  |
| MSMEG_5083 |  | Membrane | DivIVA domain-containing protein, Cell division protein | 7.76 | 4.08 |
| MSMEG_5447 | PMT | Plasma membrane | Dolichyl-phosphate-mannose--protein mannosyltransferase, part of the GT-C superfamily of glycosyltransferases, glycosyltransferase 39 family | 7.22 | 1.32 |
| MSMEG_4237 |  | Membrane | Probable conserved transmembrane protein | 5.83 | 3.43 |
| ftsH |  | Plasma membrane | ATP-dependent zinc metalloprotease FtsH, Cell division protein, Membrane-bound protease, Plays a role in the quality control of integral membrane proteins | 5.69 | 4.39 |
| MSMEG_5062 |  | Plasma membrane | Uncharacterized protein | 5.62 | 3.93 |
| MSMEG_1929 |  | Plasma membrane | Conserved alanine valine rich protein | 5.60 | 3.91 |
| MSMEG_6899 |  | Plasma membrane | Predicted to be in the GT-C superfamily of glycosyltransferases, glycosyltransferase 87 family | 5.58 | 4.12 |
| MSMEG_3155 |  | Plasma membrane, Cell wall, Extracellular region | Band 7 protein, Flotillin-like protein | 5.52 | 4.80 |
| MSMEG_5238 |  | Membrane | Uncharacterized protein | 5.40 | 3.31 |
| lspA |  | Plasma membrane | Lipoprotein signal peptidase | 5.23 | 3.40 |
| MSMEG_0977 |  | Membrane | Uncharacterized protein | 5.17 | 3.99 |
| MSMEG_0070 |  |  | Uncharacterized protein | 5.11 | 2.70 |
| MSMEG_5268 |  | Plasma membrane | Probable proline and glycine rich transmembrane protein | 5.10 | 3.06 |
| yajC |  | Plasma membrane | Sec translocon accessory complex subunit YajC | 4.93 | 4.36 |
| MSMEG_6027 |  | Plasma membrane | Major facilitator superfamily protein MFS_1 | 4.90 | 5.40 |

**Supplementary Figure 2:**
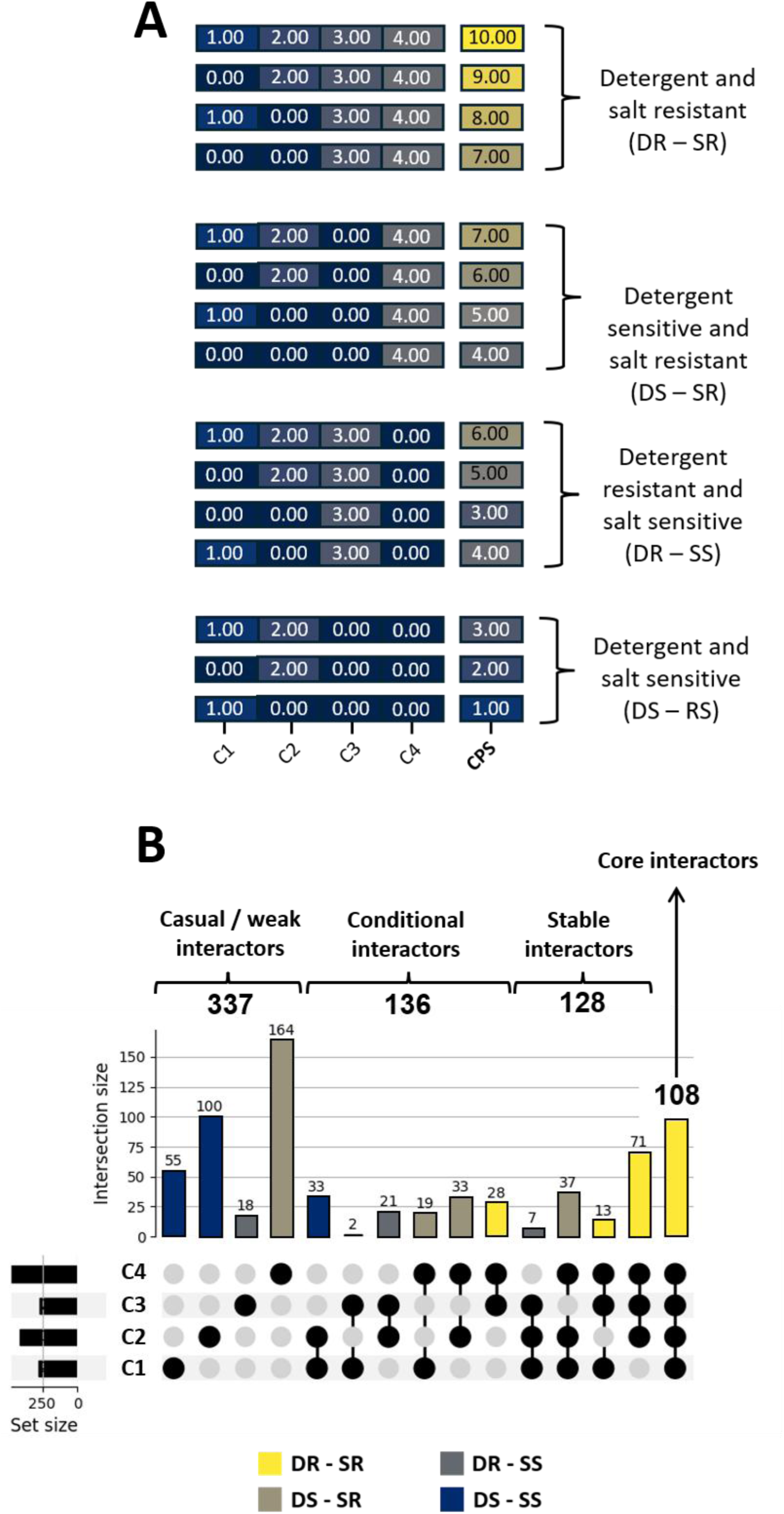
Calculation of the cumulative persistence score (CPS) and definition of biochemical resistance profiles. **(A)** Schematic representation of the cumulative persistence score (CPS) and classification of MtPMT interactors based on their resistance to biochemical perturbations. Each protein is assigned a score according to its detection across four AP–MS conditions of increasing stringency (C1–C4), with weights of 1, 2, 3, and 4 attributed to C1, C2, C3, and C4, respectively. The CPS is calculated as the sum of these values, such that proteins detected under more stringent conditions receive higher scores. Biochemical resistance profiles were defined based on the presence or absence of proteins under detergent extraction (C3) and high ionic strength conditions (C4). Proteins detected in both C3 and C4 are classified as detergent and salt resistant (DR – SR), indicating interactions stable under both membrane solubilization and elevated ionic strength. Proteins detected in C4 but not in C3 are classified as detergent sensitive and salt resistant (DS – SR), reflecting interactions that withstand high ionic strength but are disrupted or not efficiently preserved during detergent extraction, indicating that these interactions are not favored under detergent solubilization conditions. Conversely, proteins detected in C3 but not in C4 are classified as detergent resistant and salt sensitive (DR – SS), indicating stability to membrane solubilization but sensitivity to ionic strength. Proteins not detected in either C3 or C4 are classified as detergent and salt sensitive, corresponding to weak interactions lost under both perturbations. **(B)** UpSet plot of the MtPMT interaction strata illustrating the biochemical resistance profiles of proteins across the four AP–MS conditions (C1– C4). Within each stratum, the different combinations of conditions illustrate the distinct biochemical resistance profiles used to classify interactions according to their resistance to detergent solubilization and elevated ionic strength.

**Supplementary Figure 3:**
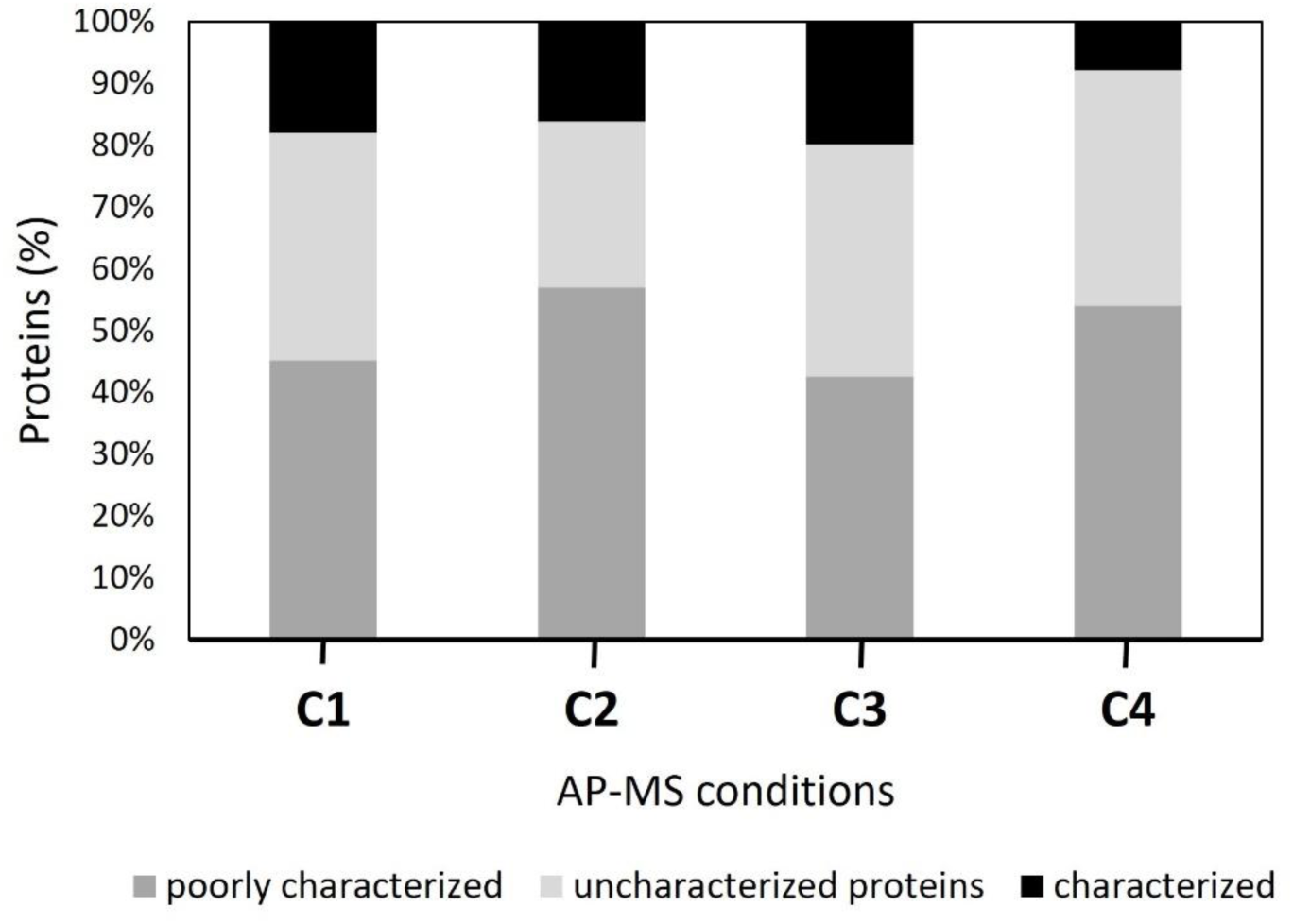
Distribution of functional annotation of MtPMT-associated proteins across AP–MS conditions. Stacked bar plots represent the proportion of uncharacterized, poorly characterized, and well-characterized proteins identified under each purification condition (C1– C4) based to Uniprot and Mycobrowser annotations. *Poorly characterized* proteins include those assigned to functional categories but lacking detailed characterization and are often identified primarily by locus tags (e.g., MSMEG_). In contrast, *uncharacterized* proteins could not be assigned to any functional category and are typically annotated as “conserved proteins”, proteins with predicted domains, or simply “uncharacterized.” Across all conditions, approximately 40– 50% of proteins are poorly characterized and 20–40% remain uncharacterized, whereas only 10– 20% are well characterized. This distribution reflects the limited functional annotation of the *M. smegmatis* mc²155 genome, particularly for genes encoding membrane proteins, which are enriched under more stringent conditions (C3–C4) and represent key MtPMT-associated partners.

**Suplementary Figure 4:**
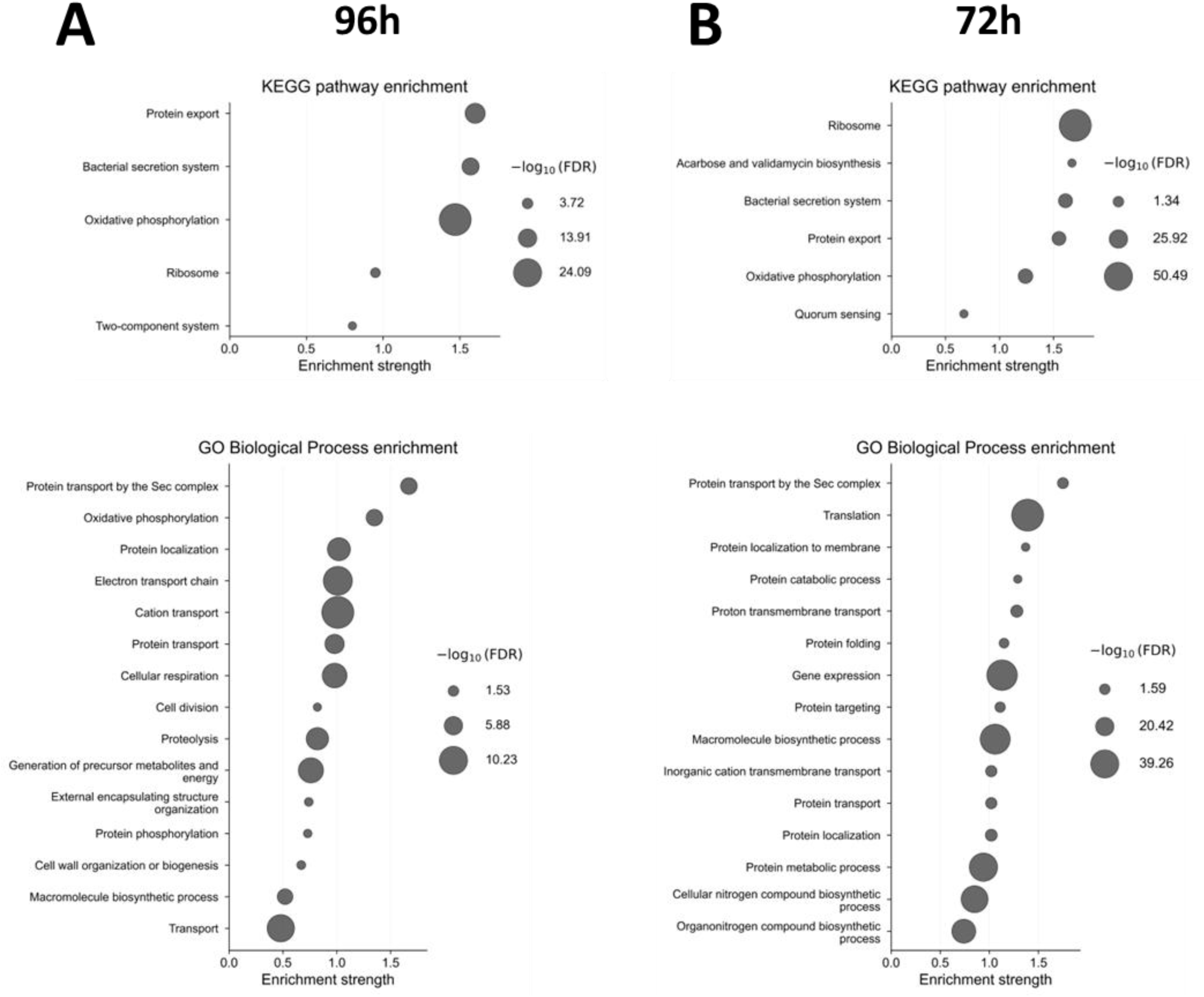
Functional enrichment analysis of MtPMT-associated proteins at 72 h and G6 h of culture. KEGG pathway and Gene Ontology (GO) Biological Process enrichment analyses of proteins identified by AP–MS at 96 h **(A)** or 72 h **(B)** of culture. Bubble plots display significantly enriched pathways and biological processes among MtPMT interactors. The x-axis indicates enrichment strength, while bubble size represents statistical significance expressed as −log₁₀(FDR). Across both conditions, enriched categories prominently include protein export and secretion pathways, oxidative phosphorylation, ribosomal components, and translation-related processes, as well as protein transport, localization, and metabolic pathways. These results support the association of MtPMT with membrane protein biogenesis, cellular metabolism, and protein homeostasis.

**Supplementary Figure 5:**
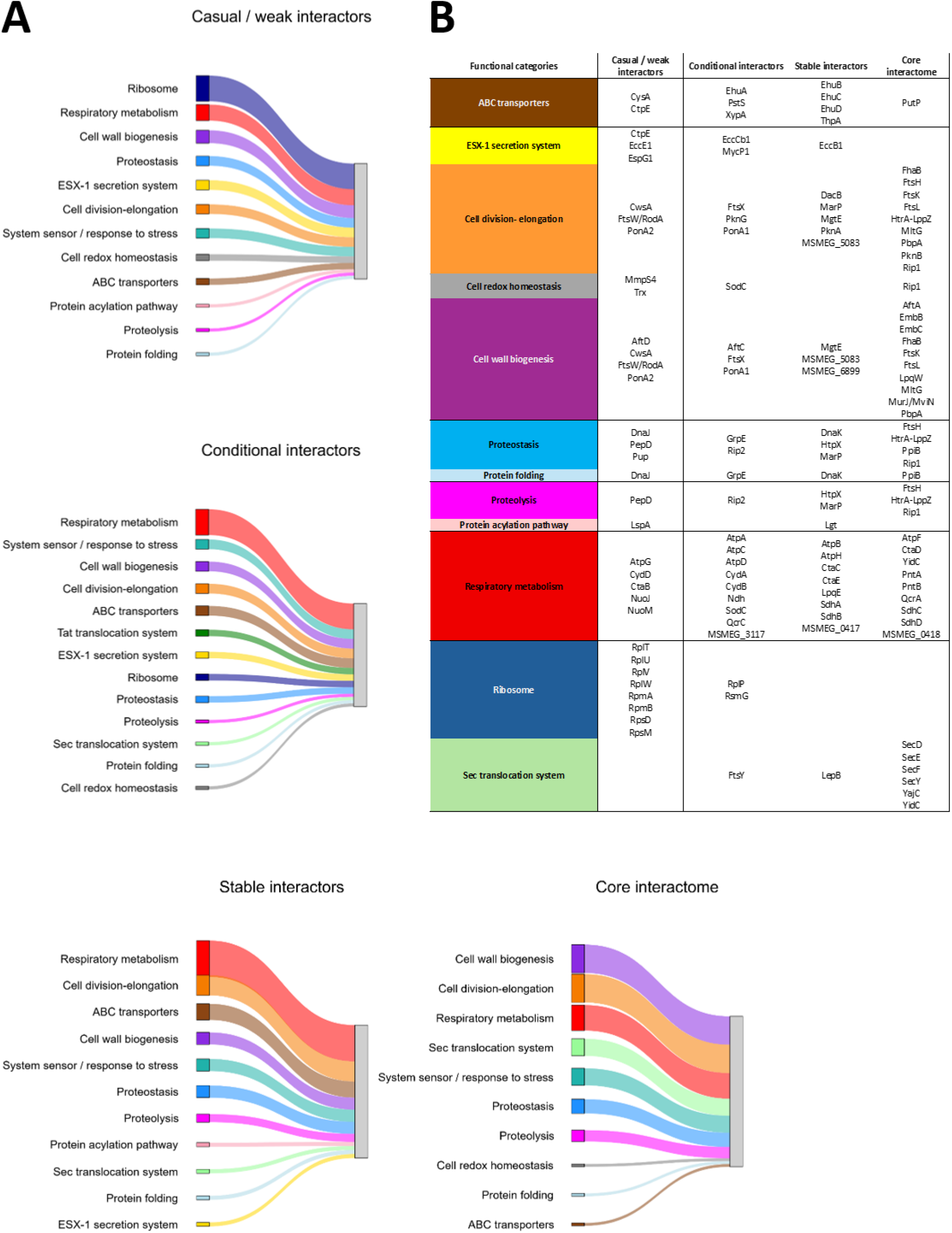
Functional classification of MtPMT interactors across interaction strata. Comprehensive list of MtPMT-associated proteins grouped according to their functional categories and interaction strata presented in Sankey diagram (**A**) and in table (**B**). Proteins are classified into casual/weak, conditional, stable, and core interactors based on their persistence across AP–MS conditions of increasing stringency (C1–C4). Functional annotation was assigned based on known or predicted biological roles (Uniprot and Mycobrowser databases), including processes such as cell wall biogenesis, proteostasis (proteolysis and protein folding), secretion systems, respiratory metabolism, and ribosome-associated functions. Only the best functionally characterized interactors are included in this analysis, and the full set of identified proteins is not exhaustively represented. **(B)** The table provides the detailed protein-level information underlying the Sankey diagram presented in **A** and in **Fig. 3B**, which summarizes the distribution of interaction strata across functional categories.

**Supplementary Figure 6:**
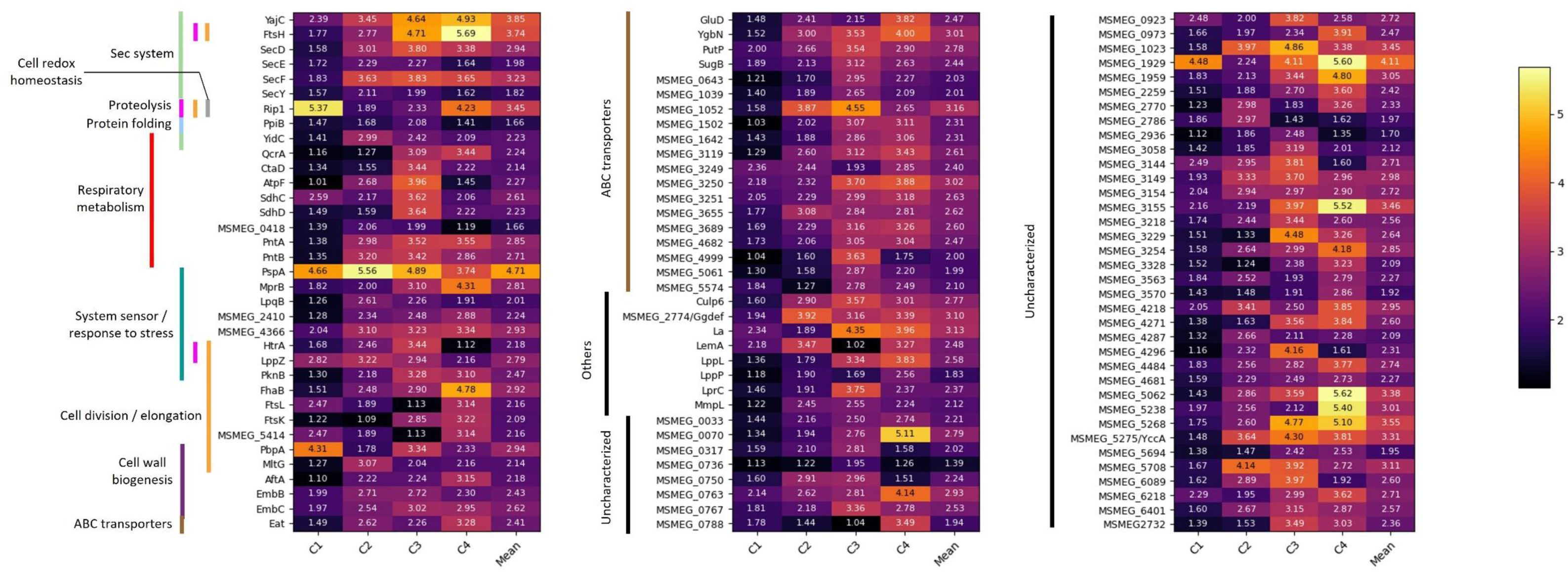
Comprehensive heatmap of the MtPMT core interactome. Heatmap representing the log₂(fold change) values of the 108 proteins constituting the MtPMT core interactome across the four AP–MS conditions (C1–C4), along with their mean enrichment values. Proteins are grouped according to functional categories, which are annotated alongside the heatmap and include Sec-dependent translocation, protein folding, proteolysis, respiratory metabolism, system sensing/stress response, cell division and elongation, cell wall biogenesis, and ABC transporters. A substantial fraction of the interactome—approximately one-thirds of the identified proteins—are functionally uncharacterized, highlighting the extent of unexplored components associated with the MtPMT membrane platform. In addition, several proteins assigned to functional categories remain poorly characterized and are identified primarily by locus tags (e.g., MSMEG_), further emphasizing the limited functional annotation of the MtPMT-associated network.

**Supplementary Figure 7:**
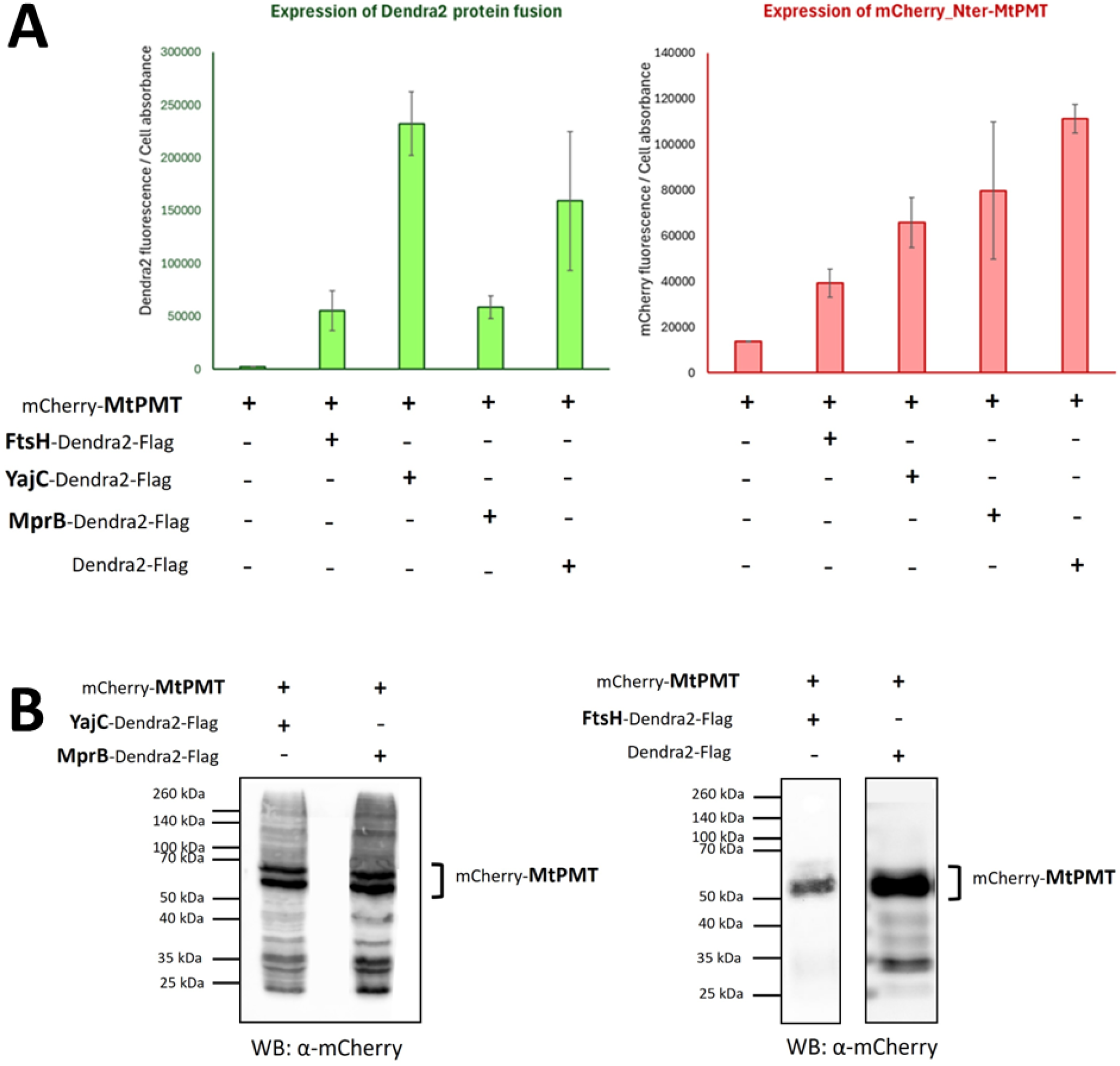
Validation of expression of fluorescent fusion constructs used for reciprocal affinity purification and FRET analyses. **(A)** Quantification of fluorescence signals in live bacteria to assess expression levels of Dendra2-tagged proteins and mCherry–MtPMT upon co-expression of the indicated constructs. Fluorescence intensity of Dendra2 (left) or mCherry (right) was measured and normalized to cellular absorbance, providing a proxy for protein expression levels *in vivo*. Data represent mean values with associated variability across replicates. **(B)** Western blot (WB) analysis of mCherry–MtPMT expression in cell lysates using an anti-mCherry antibody. Cells co-expressing mCherry–MtPMT with the indicated Dendra2-tagged proteins were analyzed to qualitatively assess protein expression. The mCherry–MtPMT fusion protein is detected at the expected molecular weight (∼50–70 kDa), confirming consistent expression across conditions. The additional bands likely correspond to aggregated or truncated forms of the protein. “+” and “–” indicate the presence or absence of the indicated constructs, respectively.

**Supplementary Figure 8:**
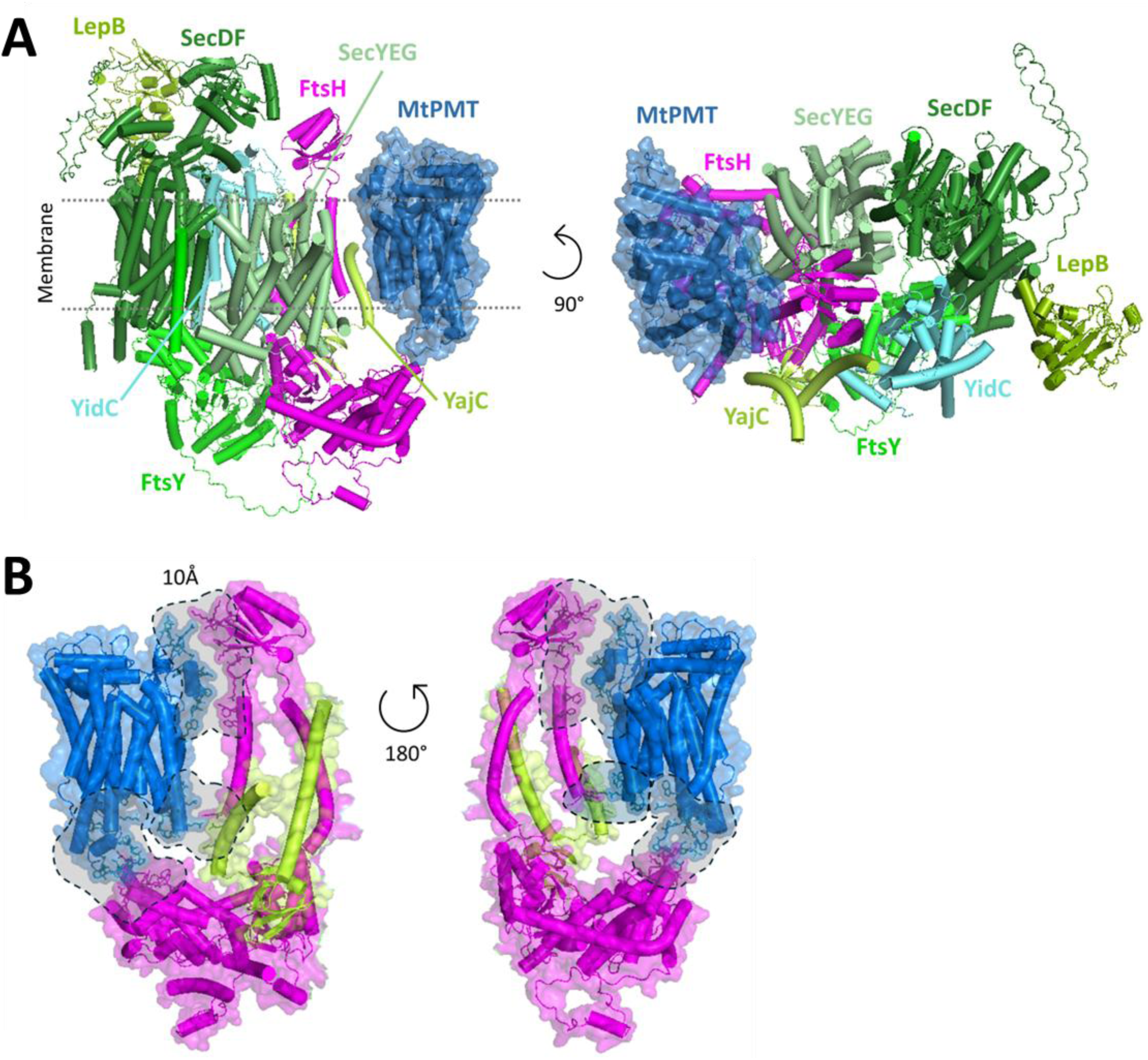
Additional structural views of the MtPMT-system Sec model and interaction interfaces. **(A)** Alternative orientations of the AlphaFold3-predicted MtPMT-containing Sec-associated complex. Proteins are represented in cartoon, with MtPMT shown in both cartoon and surface representations. The membrane is schematically indicated. These views highlight the spatial organization of MtPMT relative to Sec components, including YajC and FtsH, from different angles of a 90° rotation (side view in left, top view in right). ipTM = 0.36; pTM = 0.41 **(B)** Detailed view of the MtPMT–FtsH–YajC interaction interfaces. Proteins are represented in cartoon and semi-transparent surface. Amino acids involved in interfaces are highlighted. Regions of proximity within 10 Å are indicated by grey semi-transparent and dashed surfaces, delineating putative interaction interfaces. Two orientations (180° rotation) are shown to visualize the distribution and extent of these surfaces. The list of amino acid residues involved in these interfaces is provided in **Supplementary Table 2**.

**Supplementary Table 2:**
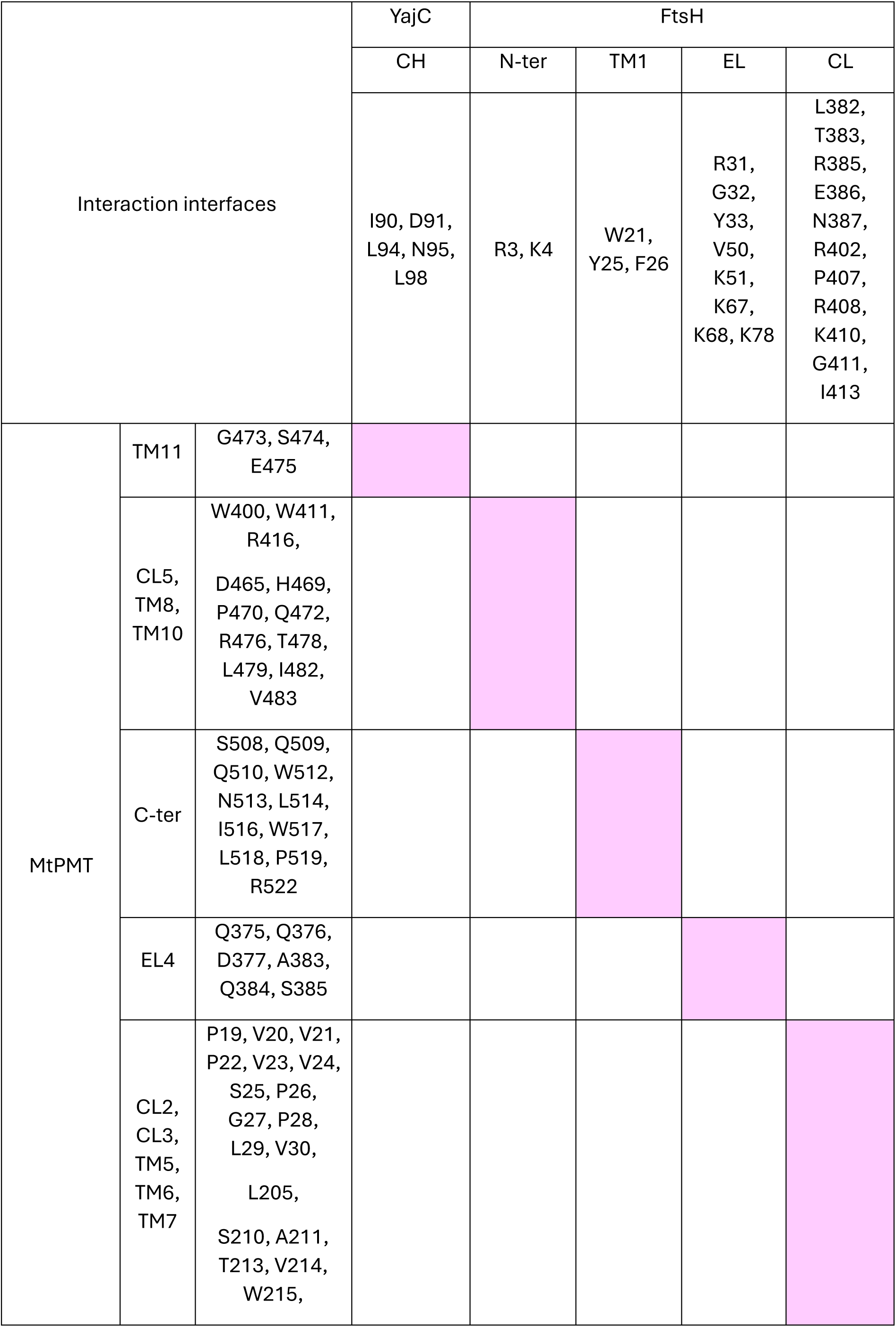
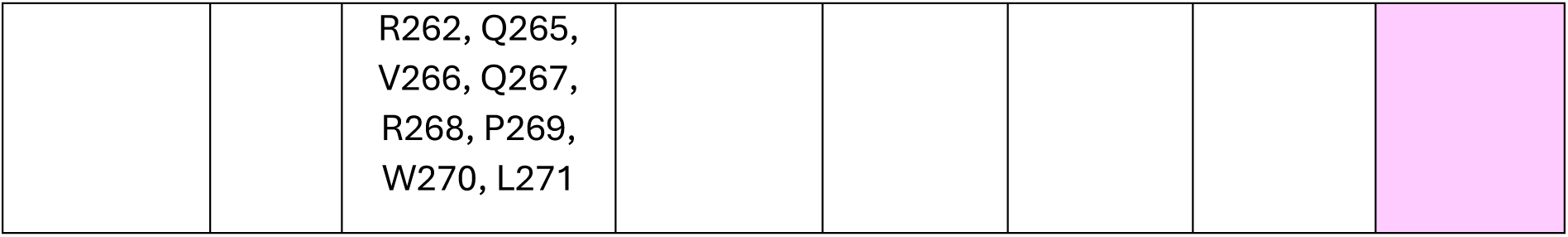
Mapping of amino acid residues involved in MtPMT interaction interfaces with YajC and FtsH. Amino acid residues participating in interaction interfaces were identified from the AlphaFold3 structural model (provided in **Fig. 5** and **Supplementary Figure 8**) and distance analysis (<10 Å) performed using PyMOL. Residues are listed according to their respective structural domains in MtPMT, YajC, and FtsH. Light pink boxes indicate detected interaction interfaces between specific domains of MtPMT and those of YajC or FtsH. Domain annotations include transmembrane helices (TM), cytoplasmic loops (CL), cytoplasmic helix (CH), and extracytoplasmic loops (EL).

**Supplementary Figure 9:**
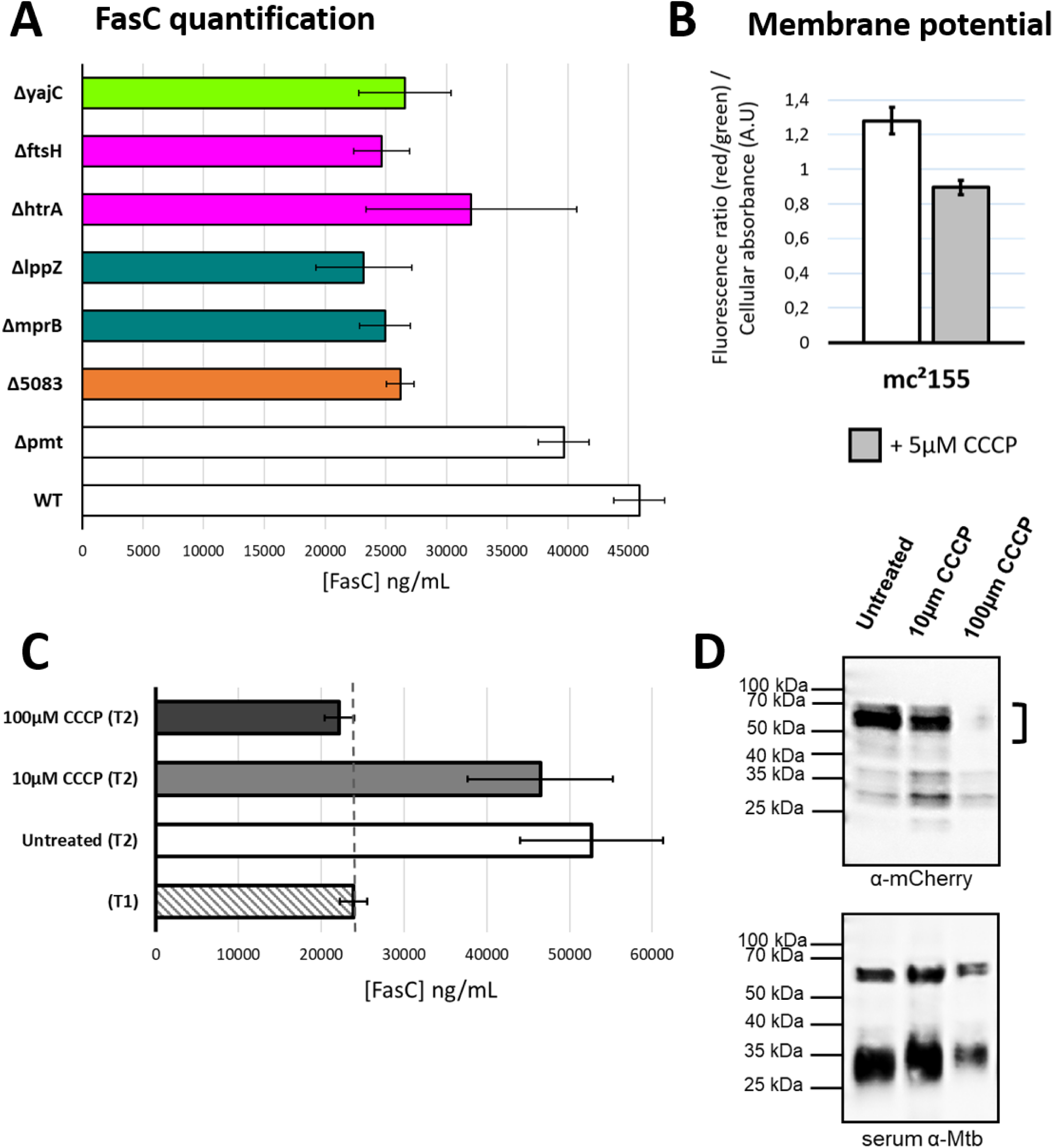
Impact of MtPMT-partners deficiency and membrane depolarization on protein expression. **(A)** Quantification of the *O*-mannosylation reporter protein FasC [1] by direct ELISA in *M. smegmatis* knockout strains. Deletions of selected MtPMT-associated partners (ΔftsH, ΔyajC, ΔmprB, ΔhtrA, ΔlppZ, Δ5083) result in reduced FasC levels compared to the wild-type (WT) and the Δpmt strains. Data are presented as mean ± s.d. of four independent biological clones, each analyzed in technical triplicates. **(B)** Membrane potential measurement. Red/green ratio of the fluorescent probe DiOC₂(3) quantify by spectrofluorimetry on alive WT strain mc²155 treated or not with 5µM of protonophore CCCP. Fluorescence intensity is normalized to cellular absorbance (arbitrary units, A.U.). CCCP induces cellular depolarization, indicated by an increase in green fluorescence relative to red one. Data are presented as mean ± s.d. of three independent biological clones, each analyzed in technical triplicates. **(C)** Effect of membrane depolarization on FasC secretion. FasC levels were quantified by ELISA before treatment (T1) and after 24 h (T2) under untreated conditions or in the presence of CCCP (10 µM or 100 µM). The dashed line indicates the FasC level at T1. FasC secretion is preserved under untreated conditions and 10 µM CCCP, but is abolished at 100 µM CCCP, indicating a strong impairment of reporter production and/or secretion under severe membrane stress. Data are presented as mean ± s.d. of three independent biological clones, each analyzed in technical triplicates. **(D)** Western blot analysis of mCherry–MtPMT expression following CCCP treatment. Cell lysates collected after 24 h treatment were probed with an anti-mCherry antibody. The bracket indicates the expected molecular weight of the mCherry–MtPMT fusion protein. Expression levels are comparable between untreated and 10 µM CCCP conditions but are markedly reduced at 100 µM CCCP. Total protein loading was controlled using a serum of whole cell lysate minus LAM *Mtb* H37Rv, confirming similar protein amounts across samples.

